# Network fixation expansion microscopy

**DOI:** 10.64898/2026.10.07.756227

**Authors:** Yixi Liu, Chi Zhang, Corban Swain, Edward S. Boyden

**Author notes:** Corresponding author (E.S.B.).

## Abstract

Expansion microscopy (ExM) physically magnifies preserved biological specimens by using a specimen-permeating hydrogel to pull biomolecules, such as proteins, apart from each other. ExM protocols exhibit a tension between the need to initially fix proteins to each other during sample preservation, to retain proteins and their nanoscale organization, and the need to separate proteins from each other isotropically, to achieve physical magnification. We here propose to address this tension by fixing proteins directly to the hydrogel network itself, which then serves as a common substrate for both initial fixation and final separation. We equip proteins with chemical handles, then form a specimen-permeating hydrogel that binds the handles, in a ∼15-minute chemical process conducted entirely at −20°C. This network fixation ExM (nfExM) procedure enables excellent epitope retention and nanoscale organization preservation, in cultured cells. nfExM may help support ultrastructural investigations of protein complexes and interactions.

## Introduction

Expansion microscopy (ExM), which physically magnifies preserved biological specimens via a chemical process, enables ordinary microscopes to perform nanoimaging^1^, and is finding widespread use throughout biology^2^, with perhaps over a thousand experimental papers and preprints to date^3^. In ExM, a sample is embedded in a densely permeating hydrogel, and swollen through the addition of water, so that biomolecules anchored to the hydrogel are separated from one another, enabling them to be better resolved on a conventional, diffraction-limited microscope. When labels that bind to specific biomolecules (e.g., fluorophore-conjugated antibodies against specific proteins) are delivered post-expansion, the decrowded state between separated biomolecules makes more room for such labels to bind, than when labels are administered pre-expansion^4–9^. ExM protocols typically involve four key steps. First, biomolecules are covalently equipped with handle-like molecules bearing a polymer-incorporable group.

Second, a hydrogel is formed (mesh size ∼1-2 nm^10^) that weaves its way throughout the specimen, inside and outside cells, and intercalating between biomolecules, covalently binding to the handles (and thus anchoring biomolecules to the polymer network). Third, the specimen is softened (typically through enzymatic, heat, and/or detergent treatment), preparing biomolecules to be separated from each other. Finally, the sample is exposed to water which causes swelling (typically by 4x-20x in a single expansion step, although the process can be iterated^7,11,12^).

Before these four steps begin, however, there is a preprocessing step, which is often taken for granted: the sample is chemically fixed, usually with covalent fixatives such as formaldehyde, which cross-link proteins to their neighbors. Such fixation is important for retaining, and preserving the nanoscale relative organization of, biomolecules such as proteins, key to the integrity of the entire process. However, this desire for strong chemical cross-linking presents a challenge, by complicating the final swelling step, during which clean, isotropic separation of biomolecules from each other is desired. Fixing molecules to each other too strongly could jeopardize the separation process, potentially yielding artifacts such as cracking, or poor protein separation. Thus, ExM protocols exhibit a tension between the first and last steps: one wants biomolecules, such as proteins, locked securely to each other at the start of the ExM procedure, but easily separable from each other at the finish.

To address this tension, we developed a protocol, network fixation expansion microscopy (nfExM), which fixes proteins to the hydrogel itself. In other words, the hydrogel serves as both the initial fixation scaffold holding everything in place, as well as the final separation actuator that pulls proteins apart from each other, thus resolving the tension between initial and final steps. But, in order for the hydrogel to serve as an initial fixation scaffold, it must be synthesized throughout the cell, and proteins must be bound to it, as quickly as possible, and under conditions that prevent degradation of cellular structure. To achieve the latter, we aimed to conduct the entire procedure at as cold temperatures as possible.

We achieved high speed, low temperature, anchoring and hydrogel formation through a series of chemical design choices. First, we administered acrolein, which equips amines with vinyl groups, to specimens, thus rapidly equipping proteins with polymer-incorporable handles. This treatment was followed by administering a low-freezing point monomer solution containing a UV-driven initiator, and then triggering the polymerization with UV light. Going from living cells to being fully polymerized took about 15 minutes; furthermore, the process was conducted entirely at −20oC. In more detail: to conduct the whole procedure at cold temperatures required several innovations: the acrolein was administered in methanol, for example, and the monomer solution was designed so that freezing point depression would keep it liquid at −20oC. Finally, the use of UV to initiate polymerization allowed the process to be completed without necessitating heating, as is commonly used to trigger polymerization in ExM. We validated nfExM by examining multiple protein targets in cultured cells, finding protein retention and nanoscale organization to be excellent, as assessed by post-expansion staining, without the expansion cracking associated with conventional strong fixation. We thus anticipate that nfExM could be of use for many scientific questions involving protein complexes and interactions.

## Results

Before arriving at the nfExM concept, we tried to see if more straightforward optimizations of ExM could resolve the seeming contradiction between strong initial fixation of proteins, and ultimate separation of proteins, with the core assessment being the quality of post-expansion staining of proteins with antibodies. These explorations were aimed at breadth, not depth, and since they generally indicated a failure to result in good post-expansion staining, we did not focus on repeating experiments to the point of statistical significance; nevertheless, we thought that it would be of use to the community to report our findings, even if they are a bit anecdotal, rather than to not report them at all.

First, starting with the basic proExM protocol (**Fig. 1A**), we asked whether we could bolster the fixation, to better retain proteins and their nanoscale organization. A strong covalent fixative like glutaraldehyde (GA) is excellent at locking biomolecules to each other, and is commonly used in the preparation of samples for classical nanoimaging methods like electron microscopy^13^. GA has been used for ExM purposes, at low concentrations (e.g. 0.1% w/v GA, mixed with 3% w/v paraformaldehyde (PFA)), followed by proteolytic digestion that requires pre-expansion staining (since such digestion destroys most epitopes, precluding post-expansion staining)^14^. We attempted to create a post-expansion staining form of ExM compatible with GA-fixed samples, by trying a variety of softening methods (e.g., sparse-cutting proteases) that would potentially spare epitopes for post-expansion staining, using microtubules in cultured cells as a testbed; our attempts failed, yielding two characteristic failure outcomes - gel cracking upon expansion, and microtubule discontinuity in post-expansion β-tubulin staining (see **Supp. Table 1** for a list of the chemicals and outcomes). Thus, combining a strong fixative with a protease was unable to balance strong initial fixation, and isotropic final separation.

**Figure 1.**
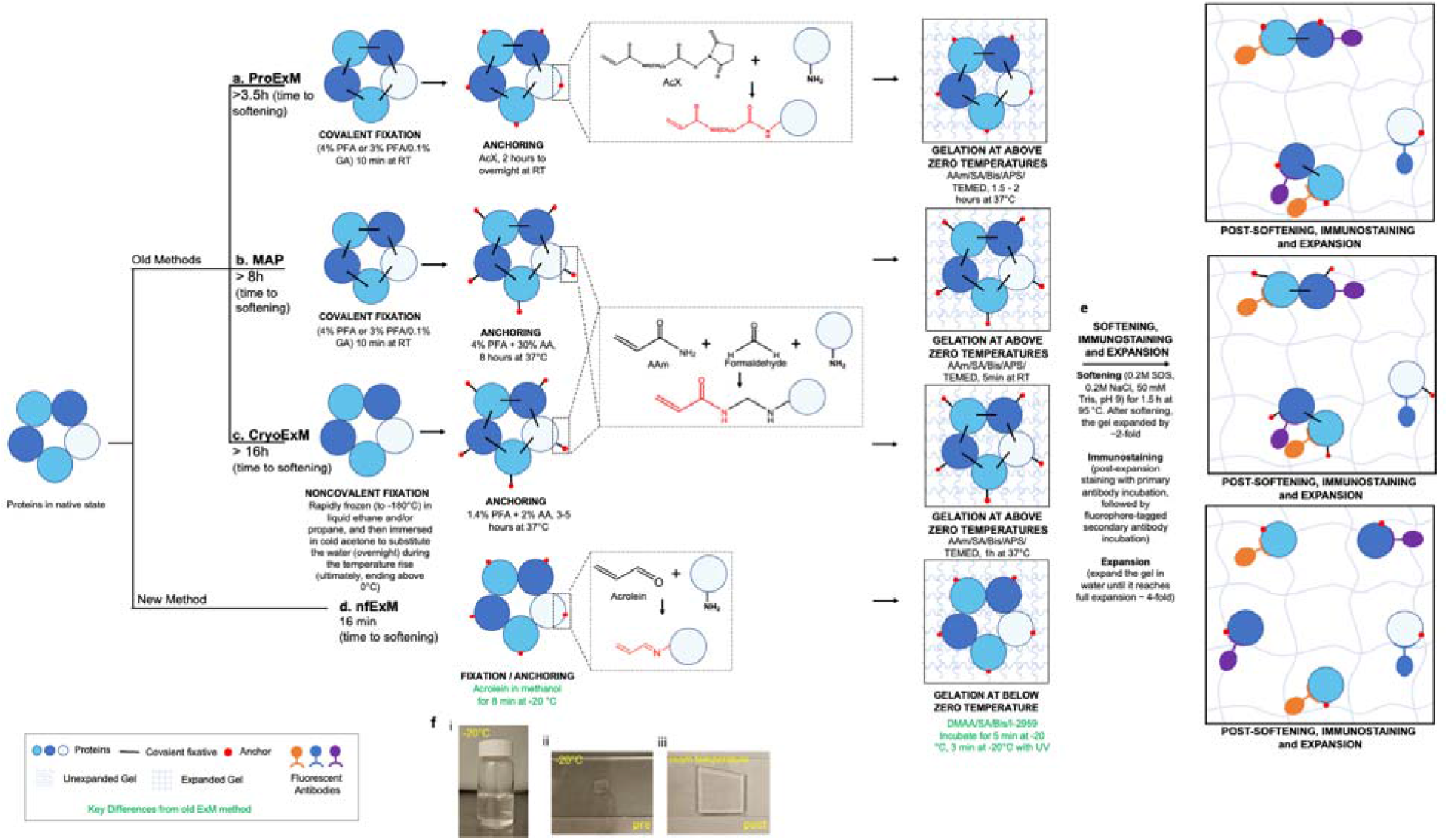
Network fixation expansion microscopy (nfExM) workflow for cultured cells. (**a–d**) Comparison of the workflow of nfExM and previous versions of ExM for cultured cells, up to the end of the gelation step. Key differences between nfExM and previous versions of ExM are highlighted in green text in d. Abbreviations: PFA, paraformaldehyde; GA, glutaraldehyde; AcX, 6-((Acryloyl)amino)hexanoic acid, succinimidyl ester; AA, acrylamide; SA, sodium acrylate; Bis, N,N′-methylenebisacrylamide; APS, Ammonium persulfate; TEMED, N,N,N′,N′-Tetramethylethylenediamine; DMAA, N,N-dimethylacrylamide; I-2959, 2-Hydroxy-4′-(2-hydroxyethoxy)-2-methylpropiophenone; SDS, sodium dodecyl sulfate; NaCl, sodium chloride; Tris, Tris(hydroxymethyl)aminomethane; RT, room temperature. (a) In proExM^8^-style ExM, cultured cells are first covalently fixed with 4% w/v PFA or a mixture of 3% w/v PFA and 0.1% w/v GA (choice determined by target structure) at RT for 10 minutes; then, proteins are equipped with a polymerizable handle by reacting them with a chemical, AcX, that binds amines and bears a vinyl group, incubating for 2 hours to overnight at RT. Next, the monomer solution (8.625% w/v SA, 2.5% w/v AA, 0.15% w/v Bis, 2M NaCl in PBS) containing 0.2% w/v APS and 0.2% w/v TEMED, is added at RT. Then, the gelation is performed at 37°C for 1.5-2 hours. (b) In MAP^5^-style ExM, cultured cells are first covalently fixed with 4% w/v PFA or a mixture of 3% w/v PFA and 0.1% w/v GA (choice determined by target structure) at RT for 10 minutes; then a mixture of PFA and AA in PBS is added at 37oC, so that the AA anchor could be coupled via PFA to the target biomolecule. This anchoring process takes 8 hours at 37°C. Next, the monomer solution (7% w/v SA, 20% w/v AA, 0.1% w/v Bis in PBS) containing 0.5% w/v APS and 0.5% w/v TEMED, is added at RT. Then, the gelation is performed at RT for 5 min. (c) In CryoExM^19^, cultured cells are first rapidly frozen (to −180oC) and then immersed in cold acetone, a noncovalent fixative, followed by raising the temperature of the acetone until it goes above 0oC, throughout which period the water is being replaced by acetone. The sample is then rehydrated at above zero temperatures, and the covalent fixative PFA along with the polymer-incorporable anchor AA is introduced at 37oC in PBS. This anchoring process takes 3-5 hours at 37°C. Next, the monomer solution (19% w/w SA, 10% w/w AA, 0.1% w/w Bis in PBS) containing 0.5% w/v APS and 0.5% w/v TEMED is added, and the cells are incubated in monomer solution on ice for 5 min. Then, the gelation is performed at 37°C for 1 hour. (d) In nfExM, cultured cells are first immersed in cold acrolein in methanol at −20°C. The aldehyde group on acrolein can equip amines on proteins with vinyl groups, so the proteins can be anchored to the polymer in a later step. Cells are then incubated in monomer solution (9.7% w/w SA, 4.9% w/w I-2959, 46.7% w/w DMAA, 0.1% w/w Bis in water) for 5 min at −20°C, followed by exposure to 365 nm UV light (0.0171 mW /mm^2^) for 3 minutes at −20°C to form the hydrogel. (e) With biomolecules secured to the gel network, samples are incubated in a softening buffer (200 mM SDS, 200 mM NaCl, and 50 mM Tris, pH 9.0) for 1.5 h at 95 °C; softened samples are then washed in PBS buffer followed by immunostaining, and then the gel is fully expanded in water. (f) Photographs illustrating (i) the monomer solution, which remains in a liquid state at −20°C; (ii) the nfExM hydrogel formed at −20°C; (iii) the expanded hydrogel (same specimen as in ii, but flipped) after immersion in water at RT.

We next asked if using a cleavable fixative would help: in principle, we could apply the cleavable fixative to form covalent cross-links of proteins to each other in the initial fixation step, and then, after proteins were safely anchored to the hydrogel, cleave the fixative during the softening step so that neighbor-neighbor cross-links were abolished. Candidate cleavable fixatives were screened both for their ability to crosslink a protein solution (assessed by PAGE electrophoresis) as well as for microtubule structure preservation; again, we encountered failure (see **Supp. Table 2** for a list of the chemicals and outcomes), with cleavable fixatives generally serving as poor crosslinkers, and not preserving microtubule structures well.

We next hypothesized that if we could fix proteins directly to the hydrogel itself, and not directly to each other, in a fashion that was rapid enough and done in as high a sample integrity state as possible, then nanoscale relative positions of proteins might be accurately preserved, and there would be no strong protein-protein crosslinking needing undoing in the softening step. In electron microscopy, to preserve molecular integrity and nanoscale relative organization, chemical fixation sometimes is conducted at cold temperatures in a process called freeze substitution, in which a frozen sample, at liquid nitrogen temperature (−196oC^15^), is exposed to an organic solvent containing a covalent crosslinking fixative (e.g., glutaraldehyde)^16,17^. The fixative then crosslinks biomolecules to each other while in a liquid state, that because it is cold, reduces protein degradation and reorganization. In the fixation style used in the pioneering MAP protocol^5^ (**Fig. 1B**), formaldehyde at 37°C crosslinks biomolecules to one another, while acrylamide (AA) enables a fraction of the formaldehyde-modified target molecules to become anchored to the subsequently formed polymer network – meaning some proteins are likely fixed to each other, some only to the polymer, and some to both. In the approach of the MAGNIFY method^18^, biomolecules are first formaldehyde-fixed at 4 °C or RT—crosslinking them to one another—then treated with a novel methacrolein-supplemented monomer solution able to combine anchoring of multiple classes of biomolecules (proteins, lipids, and nucleic acids) and gelation into a single step; still, initial biomolecular fixation could be in tension with subsequent expansion. In another pioneering study, CryoExM^19^ performed freezing and freeze substitution before ExM (**Fig. 1C**), with samples first being rapidly frozen (to −180oC) and then immersed in cold acetone, a noncovalent fixative, followed by raising the temperature of the sample above 0oC, then substituting in ethanol, and finally phosphate buffered saline (PBS), ending with the cells in liquid PBS at room temperature. Formaldehyde and acrylamide were subsequently introduced at 37°C in PBS, implementing the MAP-style fixation and anchoring scheme described above. The expansion procedure then proceeded as before. We wondered if it would be possible to administer the handle, and form the polymer, all at cold temperatures. We decided to tackle the challenge of doing all the reactions at −20oC, as an initial step towards exploring moving to even lower temperatures, but that might still be of immediate value to biologists. We also decided to focus on just proteins, for this initial study.

To realize such a protocol, we first had to find an anchoring molecule that could be delivered in a solvent that is in liquid form at −20oC, and that would effectively react with proteins at such a temperature; second, we had to design a monomer solution composition that could permeate the sample, and stay in liquid form, at −20oC; and third, we had to devise a way to trigger hydrogel formation (and anchor incorporation) without raising the temperature above −20oC. Ideally, all of these steps would occur as fast as possible – at the very least, on a timescale comparable to conventional covalent chemical fixation (i.e., 10-20 minutes)^20^.

We found that acrolein, which contains an aldehyde capable of rapidly binding amine groups (we recently suggested epoxide anchors as a multimodal anchor candidate^21^, but epoxide reactions to targets, we found, were potentially slower; see **Supp. Table 2**), and a vinyl group capable of participating in free radical polymerization, could be applied to live cultured cells in −20oC methanol, where it could attach handles to proteins. Then, we tuned the concentrations of expansion microscopy monomers to result in a solution that would not freeze at −20oC, so that the solution could evenly permeate the specimen with monomers while still cold (in contrast to earlier solutions^1,8,22^ that froze at −20oC). Finally, we adapted a strategy for triggering polymerization while holding the solution at −20oC, by using a UV light-triggered initiator that does not require heating^23,24^; traditional ammonium persulfate (APS) and tetramethylethylenediamine (TEMED) initiated polymerization requires elevated temperature (e.g., 37oC) to work. The whole process takes ∼15 min, and is conducted entirely at −20oC. The remainder of the process is very similar to previous post-expansion ExM protocols. We call this technology network fixation expansion microscopy (nfExM).

Protocol design was governed by both theoretical and practical considerations. For example, −20oC was in part chosen because traditional noncovalent fixation with methanol or acetone at subzero temperatures is often done at −20°C, and a freezer at −20°C is accessible for most biology labs. Since noncovalent methanol fixation of cultured cells at −20°C is usually done for 5–10 minutes (see protocols from Thermo Fisher Scientific^25^ and Synaptic Systems^26^), we chose 8 minutes as the initial incubation duration with 0.5% w/v acrolein in methanol. (In exploratory work, we found longer incubation periods (2h, 8h, 12h, 24h) to result in cracks across microtubules within cells, as assessed qualitatively by post-expansion staining, and that shorter treatments (3min, 5min) resulted in discontinuous microtubules, but we did not explore this systematically; we focused on 8 minutes since it seemed to work well. Similarly, we chose 0.5% w/v acrolein as the concentration because this concentration is commonly used when GA is used alone for microtubule fixation^28^, and microtubules appeared continuous by visual inspection with post-expansion staining; decreasing the concentration of acrolein to 0.1% w/v resulted in discontinuous microtubules with post-expansion staining, whereas higher concentrations (e.g., 1% and 2% w/v acrolein) resulted in inconsistent outcomes (perhaps because higher-concentration acrolein was available only with lower purity; see **Methods** section, “Optimization of Acrolein Concentration for nfExM,” for details on acrolein solution preparation and reagent sourcing).

Immediately after acrolein-methanol treatment, cells were incubated for 5 minutes in monomer solution. Although classical proExM begins polymerization, at least for cell culture, immediately after adding monomer solution, we reasoned that at −20oC diffusion could be slower, and thus would benefit from an incubation period (we also tried a 1-minute incubation, and observed no difference in microtubule preservation by eye, as assessed with post-expansion staining). We chose a monomer solution recipe (**Fig. 1d**) consisting of 46.7% w/w N,N-dimethylacrylamide (DMAA, a non-ionic ExM monomer that is liquid with a freezing point of −40°C), 9.7% w/w sodium acrylate (SA, an ionic ExM monomer that aids in expansion), and 0.1% w/w N,N’-methylene-bis-acrylamide (Bis, an ExM gel crosslinker) in water. We adjusted the ratio between DMAA and SA so that the concentration of SA would be enough to provide a final expansion factor of about 4-fold, and the concentration of DMAA would be high enough to suppress the freezing point, allowing the monomer solution to remain liquid at −20°C.

We then sought to initiate polymerization in a temperature-independent manner. Using the temperature-dependent APS/TEMED initiation system of proExM, even after a week at −20oC, no hydrogel formed. We decided to use a temperature-independent photo-initiator to trigger polymer formation with UV light. Photoinitiated gelation has previously been demonstrated in the context of ExM in the PhotoExM protocol^24^. Photoinitiator I-2959 had previously been shown to form a SA/AA/Bis gel within minutes^23^. We found it could form our DMAA/SA/Bis gel within 3 minutes (365 nm light, ∼0.0171 mW/mm^2^ at the sample). The anchoring and gelation steps took ∼15 minutes to complete. At this point, the hydrogel-cell composite was stable, and its temperature could be raised above – 20°C without loss of ultrastructural preservation. We then performed a high-temperature detergent-based denaturation, which partially expanded the sample by ∼2×. Immunostaining was performed during this partially expanded state, as in other protocols such as MAP, ExR, U-ExM, and CryoExM^4,5,7,19^. Final expansion to ∼4× was achieved by incubation of the gelled sample in an excess of deionized water. We hypothesized this would result in more isotropic separation of biomolecules (**Fig. 1e**). As we had designed, the monomer solution stayed liquid at −20oC, and gelation was possible at −20oC (**Fig. 1f).**

To evaluate the performance of nfExM, we compared nfExM with proExM and CryoExM (in their non-GA-using forms), assessing microtubule intensity and continuity using post-expansion β-tubulin staining in cultured U2OS cells. Representative images of proExM, CryoExM, and nfExM samples are shown (**Fig. 2a-c**), with intensity being higher for nfExM than the other two cases (**Fig 2d**). Continuity analysis was performed by randomly selecting microtubule segments that were not overlapped with others, and tracing them manually. We defined a gap in a microtubule when the measured, background-corrected intensity value fell below a locally adaptive threshold, set as a relative intensity threshold parameter (α) multiplied by the intensity averaged across a 7-pixel stretch flanking the pixel of interest in either direction going down the microtubule. Gap quantification at α = 0.8 (**Fig. 2e-f**) is shown for illustrative purposes, with threshold sensitivity analysis (testing robustness to parameter selection across α = 0.6–0.9) in **Fig. 2g-h**. We found that nfExM exhibited a smaller gap ratio (**Fig. 2e, 2g**) and gap size (**Fig. 2f, 2h**), compared to earlier protocols (see **Supp. Table 3** for full statistics for full statistics associated with **Fig. 2**). In the gap size distribution histogram (**Fig. 2i**) nfExM gaps fell predominantly at the 1-pixel detection limit (0.041 µm, red vertical dashed line), a floor effect indicating that, at the scales we can measure, nfExM preserves microtubule ultrastructure with fewgaps; see **Supp. Note 2** for a discussion of why our algorithm cannot resolve sub-pixel gaps. Thus, nfExM is able to preserve microtubules with high fidelity.

**Figure 2.**
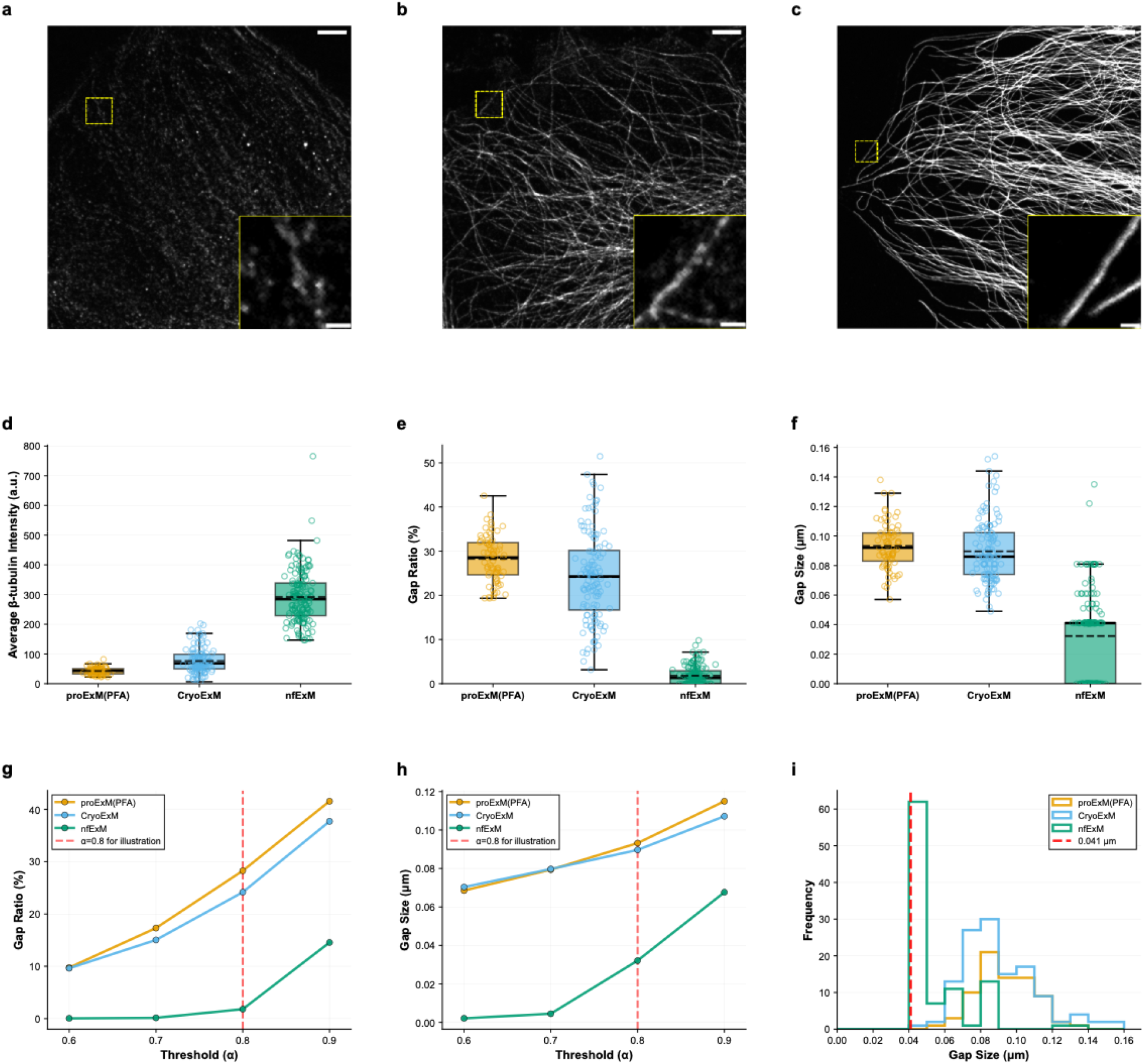
nfExM enables high-fidelity preservation of microtubules. (**a**) Representative (out of n = 10 images of 2 cultures) confocal image (maximum intensity z-projection of an image stack) of post-expansion-immunostained U2OS cells that underwent proExM (4% w/v PFA fixation) processing. The insets in (a), (b) and (c) provide a magnified view of the area indicated by the yellow dashed box in their respective images. All images were acquired under the same microscope settings. Brightness and contrast settings were adjusted for display purposes: first set by the Fiji auto-scaling function and then the minimum and maximum displayed value were set the same in (a), (b), and (c); insets share a different but uniform set of display parameters. (b) Representative (out of n = 12 images of 2 cultures) confocal image, as in a, but for CryoExM processing. (c) Representative (out of n = 10 images of 3 cultures) confocal image, as in a, but for nfExM processing. (d) Quantification of microtubule intensity in expanded U2OS cells post-proExM (PFA form), CryoExM or nfExM processing (n = 75, 124, and 155 microtubule segments from 2, 2, and 3 culture batches respectively; see **Supp. Table 3** for full statistics, including p-values). Quantification was done on raw images. Plotted is median (middle line), mean (dotted line), first and third quartiles (lower and upper box boundaries), 1.5× interquartile range (whiskers), fluorescence value averaged across all pixels of a microtubule segment (open circles). (e) Quantification of gap ratio in expanded U2OS cells post-proExM (PFA form), CryoExM or nfExM processing, for the same images analyzed in d. Plots are as in d. (f) Quantification of gap size in expanded U2OS cells post-proExM (PFA form), CryoExM or nfExM processing, for the same data analyzed in d. Plots are as in d. Gap size is in biological units (physical size divided by expansion factor of 4). (g) Threshold sensitivity analysis for gap ratio. Mean gap ratio across all traced segments plotted as a function of detection threshold parameter (α = 0.6–0.9). Red dashed line indicates α = 0.8 used for illustrative purposes in panel e. (h) Threshold sensitivity analysis for gap size. Mean gap size across all traced segments plotted as a function of detection threshold parameter (α = 0.6–0.9). Red dashed line indicates α = 0.8 used for illustrative purposes in panel f. (i) Gap size distribution for all three methods shown as step histograms. The red vertical dashed line marks the 1-pixel detection limit (0.041 µm); gaps at this limit are the minimum detectable discontinuity. Our algorithm can’t resolve gaps smaller than 1 pixel—depending on how a sub-pixel gap aligns with the pixel grid, it is either missed entirely or shows up as a 1-pixel gap (see **Supp. Note 2** for illustrative examples). nfExM shows a peak at this limit (consistent with the floor effect in f). Scale bars (in biological units used throughout this manuscript, unless otherwise noted): 2.5 µm (a, b, c), 0.5 µm (inset of a, b, c).

The nfExM monomer solution exhibited a total monomer concentration, here abbreviated T% w/v for short, that was higher than that of proExM and CryoExM, to achieve freezing point depression of the solution, but which could in principle lead to denser polymer, and more polymeric sites for protein anchors to be retained, independent of anchor identity. To investigate whether the enhanced microtubule preservation of nfExM resulted from higher T% w/v alone, we compared nfExM with a modified proExM protocol using the same high-concentration nfExM monomer solution and UV-initiated polymerization (at −20oC, as in nfExM; fixation and anchoring were done at RT, as in **Fig. 1a**) as nfExM (hereafter referred to as “high T% proExM”). If the nfExM improvement in **Fig. 2** came from higher T% w/v alone, high T% proExM should perform as well as nfExM. Microtubule intensity and continuity were assessed using post-expansion β-tubulin staining in cultured HeLa cells (**Fig. 3a-b**), with microtubule segments traced as in **Fig. 2**. nfExM exhibited a higher mean intensity (**Fig. 3c**) (n = 61 and 59 microtubule segments from 2 and 2 culture batches respectively; see **Supp. Table 4** for full statistics associated with **Fig. 3**). Gap quantification at α = 0.8 (**Fig. 3d-e**) is shown for illustrative purposes, with threshold sensitivity analysis in **Fig. 3f-g**; nfExM exhibited a smaller gap ratio and gap size compared to high T% proExM. The gap size distribution histogram (**Fig. 3h**) revealed that nfExM gaps fell predominantly at the 1-pixel detection limit (0.041 µm, red vertical dashed line), the same floor effect we saw in **Fig. 2**; in contrast, high T% proExM showed a distribution extending to larger gap sizes, indicating more and wider gaps in microtubule continuity. These experiments used the conventional temperature profiles of proExM (**Fig. 1a**); to test whether cold fixation could help achieve nfExM-level preservation, we tried cold PFA fixation (4% w/v PFA in methanol at −20°C) with both proExM and high T% proExM, but found that neither condition reached nfExM’s level of microtubule preservation (**Supp. Figure 1**).

**Figure 3.**
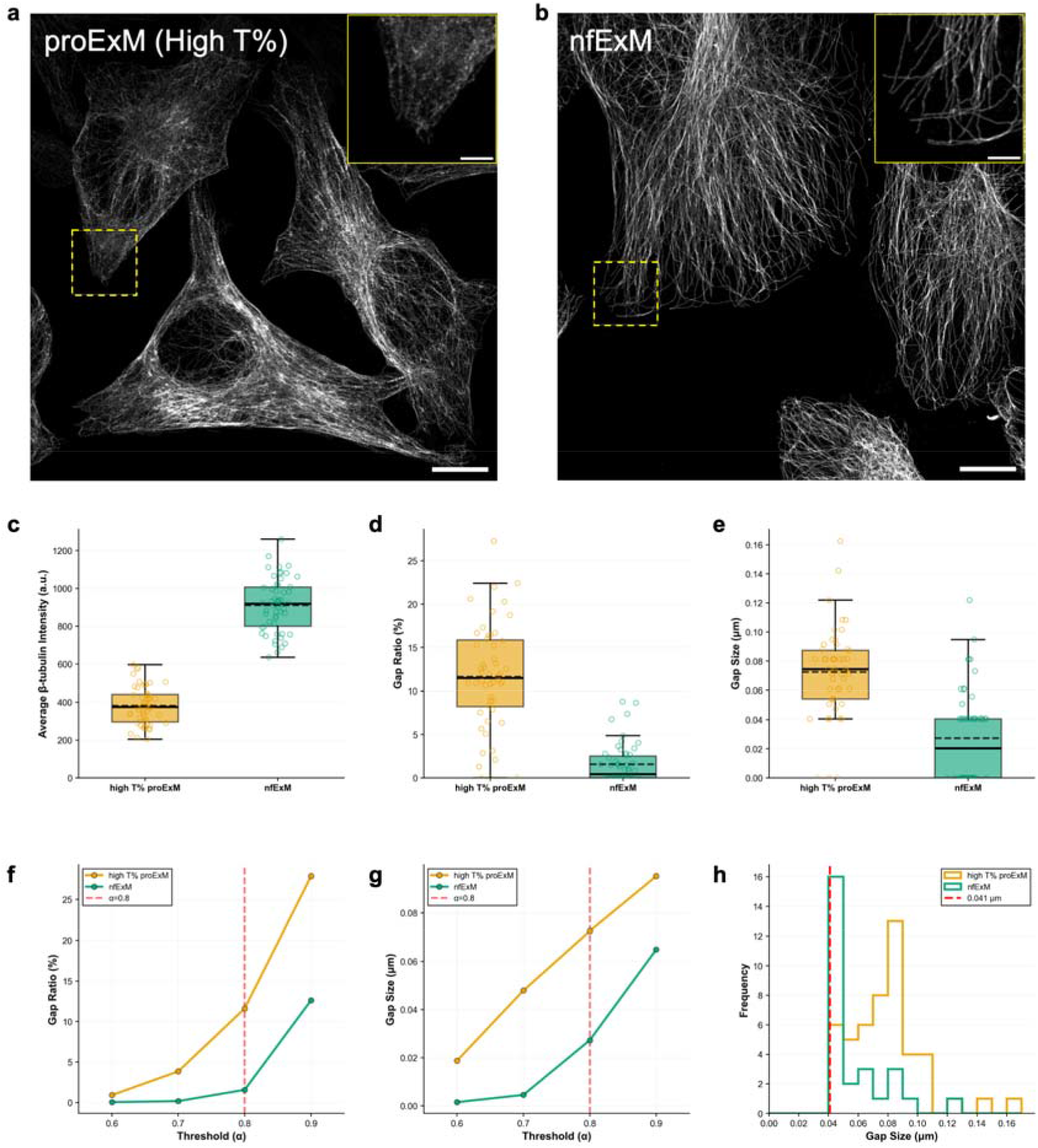
nfExM as compared to proExM with higher monomer concentration. (a) Representative (out of n = 7 images of 2 cultures) confocal image (maximum intensity z-projection of an image stack) of post-expansion-immunostained HeLa cells that underwent high T% proExM processing. The insets in (a) and (b) provide a magnified view of the area indicated by the yellow dashed boxes in their respective images. All images were acquired under the same microscope settings. Brightness and contrast settings were adjusted for display purposes: first set by the Fiji auto-scaling function and then the minimum and maximum displayed values were set the same in (a) and (b). (b) Representative (out of n = 7 images of 2 cultures) confocal image, as in (a), but for nfExM processing. (c) Quantification of microtubule intensity in expanded HeLa cells post-high T% proExM or nfExM processing (n = 59, 61 microtubule segments from 2 and 2 culture batches respectively; see **Supp. Table 4** for full statistics, including p-values). Quantification was done on raw images. Plotted is median (middle line), mean (dotted line), first and third quartiles (lower and upper box boundaries), 1.5× interquartile range (whiskers), fluorescence value averaged across all pixels of a microtubule segment (open circles). (d) Quantification of gap ratio in expanded HeLa cells post-high T% proExM or nfExM processing, for the same data analyzed in (c). Plots are as in (c). (e) Quantification of gap size in expanded HeLa cells post-high T% proExM or nfExM processing, for the same data analyzed in (c). Plots are as in (c). Gap size is in biological units (physical size divided by expansion factor of 4). (f) Threshold sensitivity analysis for gap ratio. Mean gap ratio across all traced segments plotted as a function of detection threshold parameter (α = 0.6–0.9). Red dashed line indicates α = 0.8 used for illustrative purposes in panel d. (g) Threshold sensitivity analysis as in f, but for gap size. Red dashed line indicates α = 0.8 used for illustrative purposes in panel e. (h) Gap size distribution for two methods shown as step histograms. The red vertical dashed line marks the 1-pixel detection limit (0.041 µm); gaps at this limit are the minimum detectable discontinuity. nfExM shows a peak at this limit (consistent with the floor effect in e). Scale bars: 10 µm (a, b), 2.5 µm (inset of a, b).

We next compared nfExM to a glutaraldehyde-equipped version of proExM using 3% w/v PFA/0.1% w/v GA fixation (GA form). As noted earlier, proteinase digestion (**Supp. Table 1**) resulted in microtubule discontinuity, perhaps through protein loss and epitope disruption. We accordingly tested high-heat detergent-based denaturation methods associated with past published protocols, including a denaturation-based version of proExM, dExPath, and MAP, again assessing with post-expansion β-tubulin staining in HeLa cells. Such samples frequently showed visible cracks (Fig. **4a-i**), whereas nfExM did not exhibit cracks (**Fig. 4j–m**). nfExM exhibited a higher microtubule intensity than denaturation-equipped GA forms of proExM, dExPath, and MAP (**Fig. 4n**; see **Supp. Table 5** for full statistics for **Fig. 4**), and a smaller but consistent improvement in gaps (**Fig. 4o-s**). However, ProExM, dExPath, and MAP samples exhibited gel cracking, with cracks (defined as visible discontinuities ≥1 µm in the microtubule network) present in 43.8%, 30.0%, and 43.8% of cells respectively, compared to 0% in nfExM samples (**Fig. 4t**). In sum, GA can perhaps retain more protein and reduce gaps, but at the expense of cracking (**Supp. Figure 2** presents a schematic model of why this may be so).

**Figure 4.**
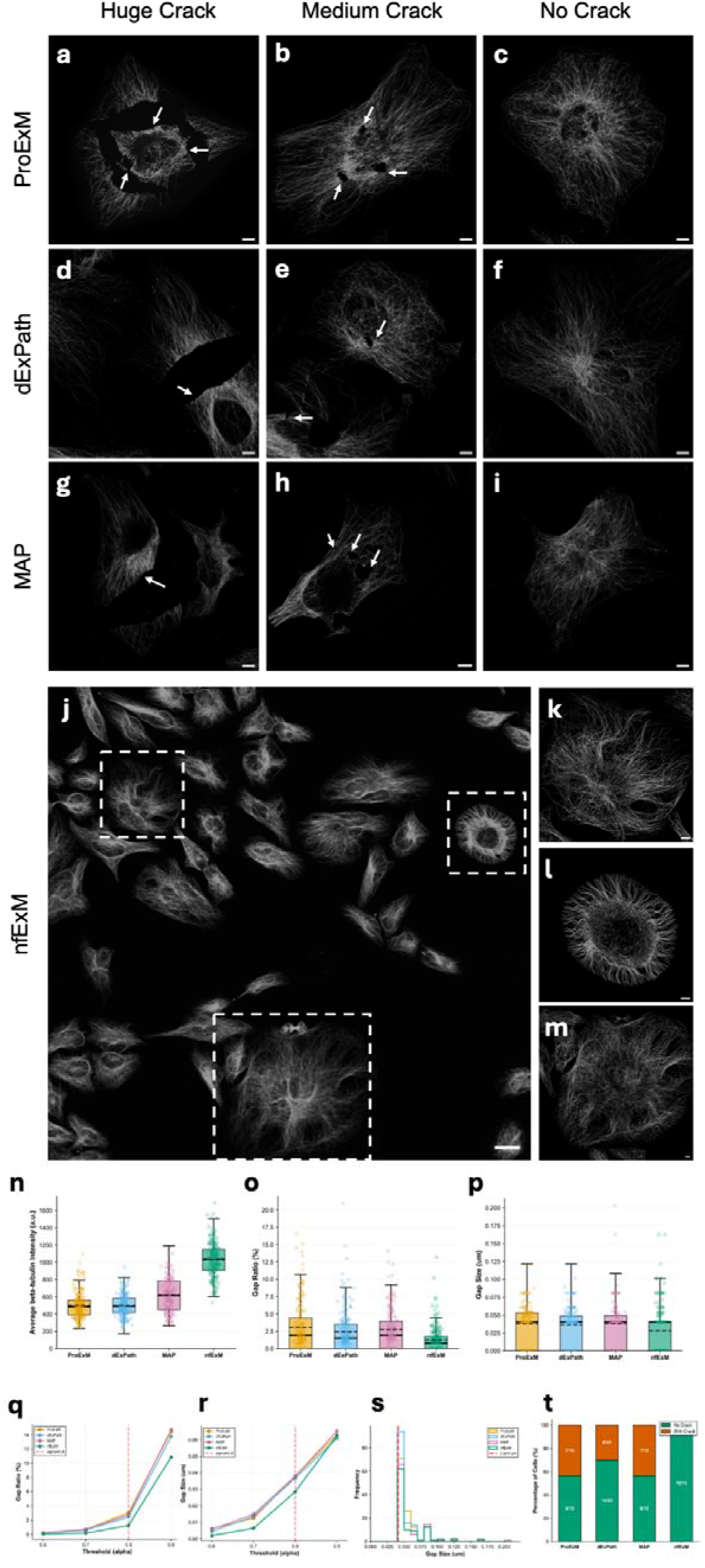
nfExM compared to glutaraldehyde-equipped forms of earlier denaturation-equipped protein-retaining ExM methods. (**a-c**) Representative (n = 16 images from 3 cultures) confocal images (maximum intensity z-projections of image stacks) of post-expansion β-tubulin immunostained HeLa cells processed with proExM (3% PFA/ 0.1%GA fix) processing followed by proExM denaturation (100 mM Tris base, 5% w/v Triton X-100, 1% w/v SDS; autoclaved at 120 °C for 1 h). Panels show examples of cells exhibiting large cracks (a), medium cracks (b), and no cracks (c). (**d-f**) Representative (n = 20 images from 3 cultures) confocal images as in (a-c), but using dExPath denaturation (20% w/v SDS, 50 mM Tris, 25 mM EDTA, 0.5% w/v Triton-X, 100 mM beta-mercaptoethanol; autoclaved at 120°C, 1 h). (**g-i**) Representative (n = 16 images from 3 cultures) confocal images as in (a-c), but using MAP denaturation (200 mM SDS, 200 mM NaCl, 50 mM Tris, pH 9.0; heated at 95°C, 1.5 h). (**j-m**) Representative (n = 16 images from 3 cultures) confocal images of HeLa cells processed by nfExM. Panel (j) shows an overview image taken with a 10x air objective lens; panels (k-m) display magnified views from the region highlighted in (j). (n) Quantification of microtubule intensity in HeLa cells processed by classical protein-retention methods of ExM with various denaturation methods and nfExM (n = 174, 193, 145, and 190 microtubule segments, respectively, from three culture batches each; see **Supp. Table 5** for full statistics, including p-values). Microtubule segments were traced only in regions without visible gel cracks. Plotted is median (middle line), mean (dotted line), first and third quartiles (lower and upper box boundaries), 1.5× interquartile range (whiskers), fluorescence value averaged across all pixels of a microtubule segment (open circles). Quantification was done on raw images. (o) Quantification of gap ratio for the same data analyzed in (n). Plots are as in (n). (p) Quantification of gap size for the same data analyzed in (n). Plots are as in (n). (q) Threshold sensitivity analysis for gap ratio. Mean gap ratio across all traced segments plotted as a function of detection threshold parameter (α = 0.6–0.9). Red dashed line indicates α = 0.8 used for illustrative purposes in panel o. (r) Threshold sensitivity analysis as in q, but for gap size. Red dashed line indicates α = 0.8 used for illustrative purposes in panel p. (s) Gap size distribution for all four methods shown as step histograms. The red vertical dashed line marks the 1-pixel detection limit (0.041 µm); gaps at this limit are the minimum detectable discontinuity. (t) Prevalence of cracking across different methods. Stacked bar chart showing the percentage of cells exhibiting visible cracks (orange) versus cells without (green) for each method. Cells were classified as “crack present” if any visible discontinuity ≥1 µm was observed in the microtubule network under confocal imaging. One cell per image (the largest by projected area) was assessed for crack presence, yielding n = 16, 20, 16, and 16 cells for proExM (GA form proExM + proExM denaturation), dExPath (GA form proExM + dExPath denaturation), MAP (GA form proExM + MAP denaturation), and nfExM, respectively, from three independent culture preparations each. All images were acquired with identical microscope settings. Brightness and contrast settings for display purpose: first set by the Fiji auto-scaling function and then manually adjusted to improve contrast for the stained structures of interest. Scale bars: 5 µm (a-i, k-m), 25 µm (j).

We next examined the performance of nfExM on the preservation of mitochondria and Golgi markers, staining for the markers TOM20 and GM130 in cultured HeLa cells. Qualitatively, mitochondrial boundaries in nfExM samples appeared more continuous than in CryoExM samples (**Fig. 5a-b**), and the Golgi apparatus signal appeared brighter in nfExM than in CryoExM (**Fig. 5c-d**).

**Figure 5.**
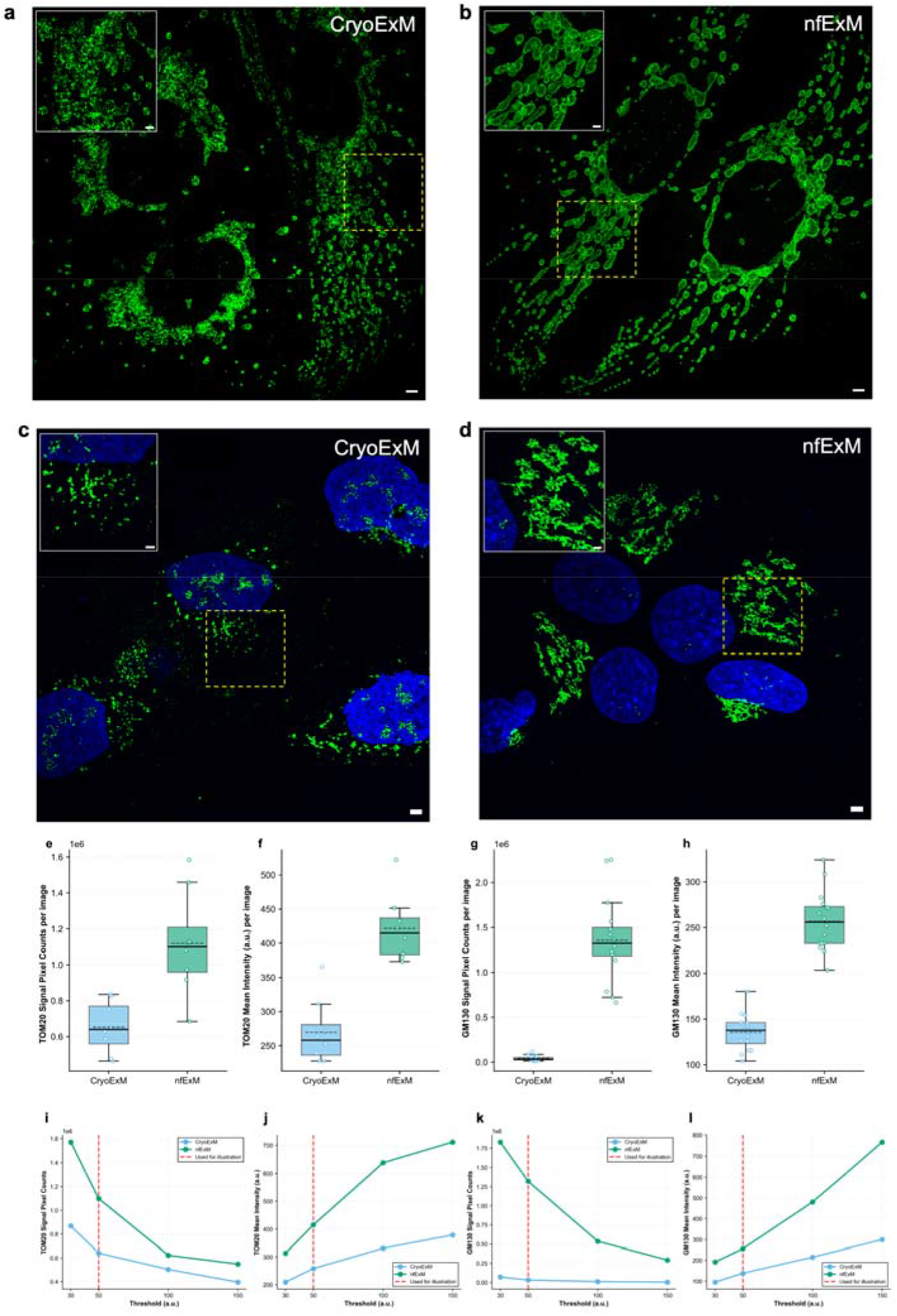
Protein retention in mitochondria and Golgi with nfExM. (**a–b**) Representative (n = 8 images from 2 cultures) confocal images (maximum intensity z-projections) of HeLa cells processed by (a) CryoExM (plunge-freezing, cold acetone fixation, anchoring and gelation at 37°C, SDS denaturation at 95°C) and (b) nfExM (acrolein-methanol anchoring and gelation at −20°C, SDS denaturation at 95°C). Cells were immunostained for the mitochondrial marker TOM20 (green). The insets in (a) and (b) show corresponding magnified views of the regions highlighted by yellow dashed lines. (**c–d**) Representative (n = 16 images from 3 culture batches each) confocal images as in (a–b), but immunostained for the Golgi marker GM130 (green), with nuclei counterstained by Hoechst (blue). The insets in (c) and (d) show corresponding magnified views of the regions highlighted in yellow dashed lines. (e) Quantification of TOM20 signal pixel counts in expanded HeLa cells post-CryoExM or nfExM processing (n = 8 images from 2 culture batches each; see **Supp. Table 6** for full statistics, including p-values). Signal pixel counts were quantified as the number of pixels with intensity above background within the segmented mitochondrial structures per image. Plotted is median (middle line), mean (dotted line), first and third quartiles (lower and upper box boundaries), 1.5× interquartile range (whiskers), and the total signal pixel count per image (open circles), summed across all segmented mitochondrial structures. (f) Quantification of TOM20 mean intensity in expanded HeLa cells post-CryoExM or nfExM processing, for the same data analyzed in e. Mean intensity was calculated as the average fluorescence intensity of all pixels within the segmented mitochondrial structures per image. Plots are as in (e), but open circles show the per-image mean intensity. (g) Quantification of GM130 signal pixel counts in expanded HeLa cells post-CryoExM or nfExM processing (n = 16 images from 3 culture batches each). Signal pixel counts were quantified as the number of pixels with intensity above background within the segmented Golgi structures per image. Plotted is median (middle line), mean (dotted line), first and third quartiles (lower and upper box boundaries), 1.5× interquartile range (whiskers), and the total signal pixel count per image (open circles), summed across all segmented Golgi structures. (h) Quantification of GM130 mean intensity in expanded HeLa cells post-CryoExM or nfExM processing, for the same data analyzed in g. Mean intensity was calculated as the average fluorescence intensity of all pixels within the segmented Golgi structures per image. Plots are as in (g), but open circles show the per-image mean intensity. (i) Threshold sensitivity analysis for TOM20 signal pixel counts, using the same images analyzed in e. Background-corrected images were segmented using four intensity thresholds (30, 50, 100, and 150 a.u.). Plotted is the mean signal pixel count across images for each method. The threshold of 50 a.u. (red dashed line) was used for illustration in panel (e-h). nfExM showed higher signal pixel counts than CryoExM across the tested thresholds. (j) Threshold sensitivity analysis as in i, but for TOM20 mean intensity, using the same images analyzed in e. Plotted is the mean of mean intensity across images for each method. (k) Threshold sensitivity analysis as in i, but for GM130 signal pixel counts, using the same images analyzed in g. Plots are as in i. (l) Threshold sensitivity analysis as in i, but for GM130 mean intensity, using the same images analyzed in g. Plots are as in j. All images were acquired with identical microscope settings and quantified from raw data. Brightness and contrast settings were adjusted for display purposes: first set by the Fiji auto-scaling function and then manually adjusted to improve contrast for the stained structures of interest. Scale bars: 2.5 µm (main images), 1.25 µm (insets).

Automated image segmentation was performed by applying a fixed intensity threshold to background-corrected images (see **Methods** for details). We found that nfExM retained more detectable structural signal (signal pixel counts), and exhibited higher mean intensity, for both TOM20 (**Fig. 5e, f**) and GM130 (**Fig. 5g,** h), compared to CryoExM (n = 8 images from 2 culture batches each for TOM20; n = 16 images from 3 culture batches each for GM130; see **Supp. Table 6** for full statistics); this conclusion was consistent across thresholds (**Fig. 5i–l**).

## Discussion

Here we describe a principle of expansion microscopy, network fixation, that may be of great utility, by enabling the expansion gel network to serve both as initial fixative and final expansion substrate. In order to create the gel network as fast as possible, and to preserve as much protein retention and organization as possible, we conducted the procedure at −20oC, going from live cells to fully polymerized in ∼15 minutes. nfExM balances two requirements in ExM that previously were in tension – the need for rapidly preserving the native nanoscale organization of proteins via high integrity initial fixation, and the need for isotropic separation of biomolecules during the final expansion step. We observed high continuity of microtubules in nfExM, and excellent continuity of mitochondrial surfaces, and high brightness of Golgi apparatus, when stained with antibodies.

Because the nfExM protocol goes from living cells to fully gelled, without an initial fixation step independent of the ExM step, we cannot perform standard measures of distortion, as done for previous ExM protocols^1,8^, where fixed cells are first imaged with a classical super-resolution microscope, and then expanded, and then re-imaged with a confocal microscope. In 20ExM^11^, microtubule width variation was used as a measure of distortion, but as of the current paper, we have not yet successfully done 20x expansion, but only 4x expansion. (When we tried iterative 20x expansion (e.g., as in ExR), we observed the gel curling after the second neutral gel re-embedding, perhaps because the nfExM gel is very dense and the re-embedding solution did not permeate it effectively; when we tried single-shot 20ExM (in the version optimized for cultured cells), we observed that the gel dissolved during high-temperature SDS denaturation for 1.5 hours at 95°C.) Future versions of nfExM that are optimized for 20x expansion could in principle be used to do microtubule sidewall analysis. That said, the polymer composition used in nfExM is related to those used in previous versions of ExM, just in different proportions, so we do not anticipate major changes in distortion. Another potential solution would involve the embedding of an intrinsic ruler, as demonstrated in the GelMap method^29^, for distortion assessment.

The nfExM protocol differs from previous protocols both in structure and in performance. Structurally, it relies on a covalent anchor being added that then is rapidly bound to a permeating hydrogel, rather than an initial fixation step followed by a later hydrogel-anchoring step. Performance-wise, it goes very quickly, with time-to-softening clocking in at 15 minutes, much shorter than previous protocols taking hours. nfExM supports post-expansion staining, which may increase the number of spatially accessible epitopes on a target protein, and increase labeling density of antibodies and their associated fluorescent signals as shown in earlier ExM protocols ^4–7,19,30^. Extending this method to a wider array of biological specimens, such as thick tissues, will require further optimization.

The requirement for UV light for initiating polymerization, for example, may require optimization for thicker specimens. In addition, our workflow requires the concurrent execution of fixation, anchoring, and polymerization at subzero temperatures without intermediate steps, which precludes immunostaining prior to polymerization. It will be of interest to see if future versions of nfExM could take place at temperatures lower than −20oC, for even better preservation.

## Methods

### Cell culture

Human bone osteosarcoma epithelial cells (U2OS; ATCC, Catalog no. HTB-96) and HeLa cells (ATCC, Catalog no. CCL-2) were cultured in Dulbecco’s Modified Eagle Medium (DMEM; Corning, Catalog no. 10013CV) with 10% v/v heat-inactivated fetal bovine serum (FBS; Thermo Fisher, Catalog no. A3840001) and 1% v/v penicillin-streptomycin (Thermo Fisher, Catalog no. 15140122) at 37 °C in a humidified 5% CO2 incubator to reach 50∼70% confluency.

### Gelation Chamber Preparation

The gelation chamber was assembled on hydrophobic glass microscope slides. To render glass slides hydrophobic for subsequent gel release, slides were immersed in a 0.2% v/v trichloro(octadecyl)silane solution (Fisher Scientific, catalog number AC147400250) in hexane, for 90 seconds at room temperature (RT). Following immersion, the slides underwent a sequential cleansing process involving a rinse with 70% v/v isopropanol at RT and subsequently with deionized water (ddH2O) at RT, to eliminate any residual silane compound or hexane. The drying process was carried out at 37°C, after which the slides were cleaned with a dry kimwipe to remove any chemical residues. Treated slides can be prepared in advance and stored at room temperature until use.

Two parallel Parafilm strips (or No. 2 coverslips) were placed ∼10 mm apart directly on the slide surface as spacers to control gel thickness. A monomer solution (27 µL for cells cultured on 12 mm diameter coverslips) was deposited at the center between the two spacers using a pipette, and the coverslip was positioned cell-side down on top of the monomer solution to close the chamber (see Supp. Fig. 3 for schematic illustration of chamber construction).

### Protease screening for post-expansion staining compatibility with GA-fixed samples (Supp. Table 1)

HeLa cells were cultured on 12 mm glass coverslips in 24-well plates. Cells were fixed with freshly prepared 3% w/v PFA, 0.1% w/v GA in PBS for 10 min at room temperature (RT), followed by reduction with 0.1% w/v sodium borohydride in 1× PBS for 7 min at RT and then quenching with 100 mM glycine in 1× PBS for 10 min at RT. After fixation, cells were washed three times with 1× PBS for 5 min each at RT. To equip proteins with polymerizable handles, cells were then incubated with 0.1 mg/mL Acryloyl-X, SE (AcX) in PBS (diluted from a 10 mg/mL stock in anhydrous DMSO) for 2h at RT without shaking.

Prior to gelation, a gelation chamber was constructed on a standard glass slide (25 × 75 mm). Spacers were created by cutting strips (∼0.3 cm wide) from the standard 4-inch (∼10 cm) width of a Parafilm M roll. Each strip was wrapped twice around the slide to form a spacer of defined thickness (∼260 µm). The spacers were positioned ∼10 mm apart to accommodate the 12 mm coverslip (see **Supp. Fig. 3** for chamber construction).

Gelation was performed using a proExM monomer solution consisting of 8.625% w/v sodium acrylate, 2.5% w/v acrylamide, 0.15% w/v N,N′-methylenebisacrylamide, and 2 M NaCl in 1× PBS. Polymerization was initiated by adding ice-cold stock solution of 10% w/v TEMED and 10% w/v APS to the ice-cold proExM monomer solution to a final concentration of 0.2% w/v TEMED and 0.2% w/v APS, with APS added last. The solution was added to the 24-well plate containing coverslips, and then incubated on ice for 3 min to permeate the specimen with monomers. The coverslip was then removed from the well and placed cell-side down onto ∼27 µL of ice-cold monomer solution (containing TEMED and APS) deposited at the center of the gelation chamber at RT. Additional solution was added from the side to fill the chamber if necessary, and the coverslip was gently pressed to ensure contact with the spacers, yielding a uniform gel thickness. Finally, the assembly was placed in a humidified box and incubated at 37°C for 1.5 h for gelation.

Following gelation, the gel was removed from the chamber, trimmed with a razor blade into an approximately 5 mm × 5 mm square shape, then one corner was diagonally removed to create an asymmetric fiducial marker for orienting the cell-containing region during imaging. The trimmed gel was transferred to a well of a 24-well plate containing the specific enzyme digestion buffer (buffer only, enzyme-free) for equilibration, 30 minutes at RT. The buffer was then replaced with fresh buffer containing the enzyme, and the gel was incubated under protease-specific conditions (see **Supp. Table 1** for buffer compositions, enzyme concentrations, temperatures, and incubation times). Following digestion, the gel was washed with PBS for 15 min followed by three washes of 30 min each. The gel was then subjected to immunostaining (at which point gels were partially expanded to ∼ 1.5-2x). First, the gel was blocked with MAXblock blocking medium (MAXblock, Active Motif, 15252) for 30–60 minutes at RT, followed by a 10 min wash with washing medium (MAXwash, Active Motif, 15254) at RT. The gel was then incubated with rabbit anti-β-tubulin primary antibody (abcam, ab6046, 1:100 dilution) in staining medium (MAXbind, Active Motif, 15253) for 5 hours at RT or 16h at 4°C. After three 30-minute washes with PBS, the gel was incubated with goat anti-rabbit Alexa Fluor 546 secondary antibody (Thermo Fisher, A-11035; 1:100 dilution) staining medium for 5 hours at RT or 16h at 4°C, followed by one wash with PBS for 30 min. Finally, the gel was expanded in deionized water for two 30-minute sessions, followed by a 16h incubation in deionized water at RT to allow for full expansion. Expanded gels were imaged promptly thereafter as prolonged storage in pure water may reduce post-expansion antibody signal due to antibody dissociation in low-ionic-strength conditions^31^.

To evaluate gel integrity, phenotypes were defined as follows: “Gel cracking” refers to >1 μm-scale (physical units) fractures of the hydrogel observed in cell-containing samples, where the gel exhibits visible fractures that cut across the cell and abruptly truncate or displace microtubule bundles at the crack planes. “Microtubule discontinuity” refers to fragmentation of the microtubule labeling into short segments or puncta with frequent, e.g. ∼150-500 nm (physical units; the lower limit is based on what is detectable using the pixel gap method, see Fig. 2i, 3h, 4s, and Supp. Fig. 4) gaps along the filaments, in contrast to continuous microtubules that can be traced for long distances.

All image scale bars in **Supp. Table 1** are 50 μm (in physical units).

Protocol references in **Supp. Table 1** are A: https://www.neb.com/en-us/protocols/2016/09/14/in-gel-digestion-protocol-for-endoproteinase-lysc-p8109; B: https://www.promega.com/resources/protocols/product-information-sheets/n/gluc-sequencing-grade-protocol/; C: https://www.promega.com/products/mass-spectrometry/proteases-and-surfactants/rlys-c-mass-spec-grade/; D: https://www.promega.com/products/mass-spectrometry/proteases-and-surfactants/arg_c_-sequencing-grade/; E: https://www.promega.com/products/mass-spectrometry/trypsin/trypsin-lys-c-mix-mass-spec-grade?catNum=V5071#protocols; F: https://www.promega.com/products/mass-spectrometry/proteases-and-surfactants/chymotrypsin_-sequencing-grade/.

### Cleavable fixative screening (Supp. Table 2)

**Design Principles.** We tried both custom-designed and commercially available cleavable crosslinking fixatives, guided by the following principles: first, the fixative molecule should be small enough to rapidly diffuse throughout the sample; second, the fixative should have multiple reactive groups that target functional groups on proteins (to achieve crosslinking); third, the fixative should be fast-acting to quickly lock biomolecules in place; and fourth, the crosslink should be cleavable. For the fixatives tested, this cleavability is afforded in one of three ways: (1) the cleavable bond is within the crosslinker itself and can later be broken by a cleavage reaction; (2) the cleavable bond is formed post-crosslinking and can later be broken by a cleavage reaction; or (3) the cleavable bond is formed at the conjugation between the crosslinker and the protein rather than in the crosslinker backbone itself.

**In Vitro Screening.** *First crosslinker choices:* At the beginning of the project, we first attempted to custom-synthesize a homofunctional traceless fixative (bi-APB; a molecule with boronic acid on both ends – see **Supp. Table 2**). Bi-APB forms protein crosslinks based on the formation of stable iminoboronates on both ends, which can be reversed by adding fructose, dopamine, or glutathione^32^, leaving no trace on the proteins – hence, “traceless.” However, our mass spectrometry results showed no desired product formation. This may be because the dissociation constant *K_d_* was too large, resulting in weak binding between proteins and this crosslinker^33^. We also analyzed this crosslinker by reacting it with protein and using an SDS-PAGE gel to assess the crosslinking degree of the reaction mixture by molecular weight (MW). After electrophoretic separation, the gels were imaged and quantified in Fiji/ImageJ using the gel analyzer tool to determine crosslinking degree. We categorized the imaged bands into two groups: first, the source band (i.e. the lowest MW band assuming no cleavage) and, second, the crosslinked bands (i.e. all bands of MW higher than the source band). Crosslinking degree (range: 0–100%) was computed as the summed integrated intensity of crosslinked bands divided by the summed intensity of all bands (crosslinked + source).

For these experiments, we desired to simulate the high protein density found in vivo (e.g., a typical protein concentration might be 20%-30% of cell volume in many cells^34^); thus, we tried to use a protein concentration of 100 mg/mL^35^. Specifically, to assess crosslinking degree for these early crosslinkers, we reacted 100 mg/mL BSA with 15 mM crosslinker in 50 mM phosphate buffer for 12 hr without shaking. Bi-APB still showed no crosslinking with these PAGE-gel based experiments.

*Second set of crosslinkers:* We then moved on to test cleavable crosslinkers, the rationale being that after protein crosslinking, we could cleave the cleavable bond in the crosslinker so that it would not prevent expansion in the later expansion step. We next tested 15 commercially-available crosslinkers. These crosslinkers were assessed for crosslinking degree by reacting with protein followed by gel electrophoresis using the same conditions as Bi-APB experiments: 100 mg/mL BSA with 15 mM crosslinker in 50 mM phosphate buffer for 12 hr without shaking.

*Third set of crosslinkers:* We also tested 7 custom-designed crosslinker candidates. For these crosslinkers, at the beginning of our design (see **Design Principles**), we had glutaraldehyde in mind but with a cleavable bond—and we wanted to compare to glutaraldehyde’s crosslinking ability (both crosslinking degree and reaction speed).

We aimed to use the same crosslinking degree assessment conditions as the first two sets of crosslinker tests; however, when we reacted 1.1% w/v crosslinker sulfonyl-bis-acetaldehyde with 100 mg/mL protein, the entire reaction mixture formed into a white, solid-like mass. The reaction mixture’s consistency was such that it could not be pipetted and run on an SDS-PAGE gel. In light of this result, we adjusted the protein concentration for this third set of crosslinkers to 10 mg/mL. Specifically, we reacted 10 mg/mL α-Lactalbumin with 1.1% w/v crosslinker in 1× PBS for 1 hr without shaking.

A few notes on the differences between the first two sets, and the third set of crosslinkers, regarding assessment conditions: **Reaction Time:** We initially set the reaction time to 1 hr considering the timescale of conventional chemical covalent fixation using PFA or GA is about 10 minutes, but the specific choice of 1 hr was arbitrary. Additionally, if a reaction takes longer than 1 hr for a cultured cell, we reasoned that we would not want such a slow-reacting fixative, as it would not be quick enough to lock biomolecules in place and biomolecules might rearrange during this time. However, when we reacted first and second-set crosslinkers with protein, the crosslinking degree was very low (<10%), so we extended the reaction time to 12 hr for this subset of crosslinkers to better discriminate between their crosslinking efficacies, purely for chemical exploration. **Protein:** We initially utilized BSA for first and second-set tests as it is a standard laboratory protein product; however, we switched to α-Lactalbumin for third-set tests because the protein purity of that reagent was higher. **Reaction Buffer:** In initial tests with first and second-set crosslinkers, we used phosphate buffer without other salts because the dissolved salts might promote excessive protein precipitation when conducting the experiments. For third-set tests, the protein concentration was lower, and we desired to test in conditions as similar as possible to cell culture experiments, so PBS (phosphate buffer with salt) was used. **Crosslinker Concentration:** The first and second-set crosslinkers were primarily in a powder form so we used molar concentration (15 mM), third-set crosslinkers were primarily in a liquid form so we used mass fraction (1.1 % w/v).

For these reasons, the stated crosslinking degree values for 1^st^/2^nd^ set crosslinkers should not be compared to third-set crosslinkers. Furthermore, we ultimately found that the *in vitro* PAGE-based crosslinking degree assessments did not correlate well to the results observed when crosslinkers were evaluated in cell culture (all of which gave poor results in cells).

**In Situ Screening on HeLa Cells.** Next, we applied cleavable fixative candidates to cultured HeLa cells. For fixation-only experiments (no ExM processing and no cleavage), cells were fixed under candidate-specific conditions, as indicated below each corresponding confocal image. Fixation-only immunostaining was performed as follows: cells were incubated in 200 μL of MAXblock blocking medium for 15 min at RT, followed by a single wash with MAXwash™ washing medium for 10 min at RT. For primary antibody staining, rabbit anti-tubulin (Abcam, catalog no. ab6046) was diluted 1:100 in MAXbind™ staining medium. Approximately 200 μL of the primary antibody solution was added to each well of a 24-well plate, followed by incubation for 2 h at room temperature. After primary antibody incubation, cells were washed three times with 1×PBS for 10 min each at room temperature. For secondary antibody staining, goat anti-rabbit Alexa Fluor 546 (Thermo Fisher Scientific, catalog no.

A-11035) was diluted 1:100 in MAXbind™ staining medium. Approximately 200 μL of the secondary antibody solution was added to each well, followed by incubation for 2 h at room temperature. Cells were washed three times with 1×PBS for 10 min each at room temperature. Microtubule preservation was assessed qualitatively from confocal images by eye. Confocal overview images were acquired either using a 40× water-immersion objective, or a 10× air objective as indicated below the confocal images. In all cases, the lower panel shows a magnified crop from the corresponding upper image.

For crosslinkers used as fixatives followed by ExM processing, cells were gently rinsed twice with prewarmed (37°C) DPBS to remove any residual cell culture media. A fixative supplemented with 1% PFA or 3%PFA in PBS was then added (note: we initially tested 1% PFA as a starting point (the choice of 1% was arbitrary). However, for some fixatives, even supplemented with 1% PFA the preservation was still very poor (similar to the condition without PFA), so in some cases we increased the PFA concentration to 3%, indicated in **Supp. Table 2**), and cells were incubated for 8 hours at RT. Cells were subsequently washed three times with PBS, 5 minutes each at RT. After removing the PBS, cells were incubated in 0.1 mg/mL AcX in PBS for 2-3 hours at RT without shaking. After anchoring with AcX, the cells were washed three times with 1x PBS, 5 minutes each.

Gelation was performed using a proExM monomer solution consisting of 8.625% w/v sodium acrylate, 2.5% w/v acrylamide, 0.15% w/v N,N′-methylenebisacrylamide, and 2 M NaCl in 1× PBS. Polymerization was initiated by adding ice-cold stock solution of 10% w/v TEMED and 10% w/v APS to the ice-cold proExM monomer solution to a final concentration of 0.2% w/v TEMED and 0.2% w/v APS, with APS added last. The solution was added to the 24-well plate containing coverslips, and then incubated on ice for 3 min to permeate the specimen with monomers. The coverslip was then removed from the well and placed cell-side down onto ∼27 µL of ice-cold monomer solution (containing TEMED and APS) deposited at the center of the gelation chamber at RT. Additional solution was added from the side to fill the chamber if necessary, and the coverslip was gently pressed to ensure contact with the spacers, yielding a uniform gel thickness. Finally, the assembly was placed in a humidified box and incubated at 37°C for 1.5 h for gelation.

After the gelation process, samples were subjected to specific softening and cleavage protocols based on the fixative chemistry. Unless otherwise specified, all incubation steps involving shaking were performed on an orbital shaker at 50 rpm.

For fixatives with a sulfone bond, gels were incubated in a solution containing 8 M urea and 50 mM dithiothreitol (DTT) in water (adjusted to pH 9.3 with NaOH) for 20 minutes at RT with shaking. This incubation accomplished softening (via urea and DTT) and crosslinker cleavage (via high pH) in the same step. While standard protocols for sulfone-based crosslinkers utilize high alkaline conditions^36^, we performed the reaction at a reduced pH of 9.3. This condition yielded sufficient cleavage to allow for expansion and might mitigate the risk of peptide bond cleavage associated with extreme alkalinity^37^.

For fixatives with disulfide bond, gels were incubated in a solution consisting of 8M urea and 50 mM DTT in water (with no pH measurement or adjustment) for 20 minutes at RT with shaking. Since the crosslinkers’ cleavable site is a disulfide moiety, the cleavage and softening actions are accomplished simultaneously by the urea+DTT solution.

For fixatives with diol bonds, gels were softened by incubation in a solution consisting of 8 M urea and 50 mM DTT in water (with no pH adjustment) for 20 minutes at RT with shaking, followed by three washes with 1x PBS for 30 minutes each at RT. The cleavage was then performed using a protocol adapted from^38^. Briefly, gels were incubated with 15 mM sodium periodate in 20 mM sodium phosphate buffer (pH 6.5) and 1M NaCl, for 30 min at RT, protected from light. The solution was then replaced with fresh 20 mM sodium phosphate buffer containing 15 mM sodium periodate, and incubation proceeded for an additional 4 hours protected from light.

For fixatives that bear two ester sites, gels were softened by incubation in a solution consisting of 8 M urea and 50 mM DTT in water (with no pH adjustment) for 20 minutes at RT with shaking, followed by three washes with 1x PBS for 30 minutes each. Cleavage was then performed by incubating gels with 0.5M hydroxylamine in 0.5M potassium phosphate buffer (pH 8.5) and 1M NaCl, incubating for 6 hr at 37 °C with shaking.

Following the specific cleavage step, all gels were washed three times with 1× PBS for 30 minutes each at RT.

For immunostaining, the gel was first incubated in 400 μL of MAXblock blocking medium for 30 minutes to 1 hour. This was followed by a single wash with MAXwash™ washing medium for 10 minutes to prepare the gel for antibody incubation. For primary antibody staining, rabbit anti-tubulin (abcam, catalog no. ab6046) was diluted 1:100 in MAXbind™ staining medium. Approximately 250 μL of the primary antibody solution was added to each well of a 24-well plate, and the gel was incubated for 5 hours at RT or 16h at 4°C. After primary antibody binding, the gel was washed three times with 1x PBS, 30 minutes each at RT. For secondary antibody staining, goat anti-rabbit AF546 (Thermo Fisher, catalog no. A-11035) was diluted 1:100 in MAXbind™ staining medium. Similar to the primary antibody step, 250 μL of the secondary antibody solution was added to each well, followed by incubation for 5 hours at RT or 16h at 4°C. The gel was washed three times with 1x PBS, 30 minutes each at RT. Finally, the gel was expanded in deionized water for two 30-minute sessions, followed by a 16h incubation at RT. The microtubules and gels were examined in confocal images by qualitative inspection, checking for cracking and microtubule continuity.

### Reagents and nfExM Monomer Solution Preparation

Sodium acrylate (SA; Pfaltz & Bauer, Catalog no. S03880) served as the ionic monomer which is critical for the gel’s expansion. N,N-Dimethylacrylamide (DMAA, Acros Organics, Catalog no. 432021000, density = 0.962 g/mL at 25°C) was employed as a co-monomer. N,N’-Methylenebisacrylamide (Bis, Sigma-

Aldrich, Catalog no. 146072) was used as a crosslinker, prepared as a 2% w/w stock solution in ultrapure water and stored at 4°C. Irgacure 2959 (I-2959, Sigma-Aldrich, Catalog no. 410896) was chosen as the photoinitiator to initiate the polymerization under UV light without requiring thermal activation.

The nfExM monomer solution was prepared as follows. First, 500 mg of I-2959 was weighed and dissolved in 5 mL of DMAA (density 0.962 g/mL at 25°C) by vortexing (Solution A); I-2959 is poorly soluble in water but dissolves readily in DMAA. Separately, 1 g of sodium acrylate was dissolved in 3.5 mL of ultrapure water (Thermo Fisher, Catalog no. 10977015), followed by addition of 500 µL of 2% w/w Bis-acrylamide stock solution (Solution B). Solutions A and B were then combined and mixed by vigorously vortexing, followed by brief sonication in an ice-water bath (approximately 1–2 minutes) in the dark to remove air bubbles introduced during vortexing. The Bis-acrylamide stock solution was prepared by dissolving 200 mg of Bis-acrylamide in 10 mL of ultrapure water and stored at 4°C. This protocol yields a final composition of approximately 9.7% w/w SA, 4.9% w/w I-2959, 46.7% w/w DMAA, and 0.1% w/w Bis (total mass ≈ 10.31 g). The prepared gelation solution can be stored at −20 °C, protected from light, for future use. The long-term stability of this gelation solution at −20 °C has not been systematically tested. However, we have used the solution stored at this temperature for two weeks, observing no notable degradation in the quality of the results, as evidenced by consistent intensity and continuity of microtubules.

### Optimization of Acrolein Concentration for nfExM

To evaluate the effect of acrolein concentration on structural preservation in nfExM, we tested varying concentrations of acrolein (0.5%, 1%, and 2% w/v). Acrolein at 0.5% w/v in methanol was purchased from Restek (Restek, Catalog no. 30645; 5000 µg/mL in methanol, equivalent to 0.5% w/v). However, since

Restek supplies a maximum concentration of 0.5% w/v, higher concentrations (1% and 2% w/v) were prepared from acrolein sourced from Oakwood Chemical (Oakwood Chemical, Catalog no. 103755-5g), which provides acrolein at a concentration of 90% (w/w). 1% or 2% w/v acrolein solutions were diluted from the 90% w/w acrolein using methanol (Thermo Scientific, Catalog no. AC326951000) and prepared freshly prior to use; the required volume of acrolein-methanol solution was transferred from the original glass ampoule into an amber microcentrifuge tube ∼1 h before the experiment, and chilled at −20°C in a metal cold block until use. We used methanol instead of acetone because the acrolein from Restek was supplied in methanol; in principle, acetone could also be used.

HeLa cells were incubated in the various acrolein solutions for 8 min at –20 °C (note: after adding the acrolein, the well was sealed with laboratory tape or adhesive plate sealing film to prevent acrolein evaporation, see **Supp. Note 1** for details of acrolein handling), followed by incubating in nfExM monomer solution for 5 min at −20C, and then followed by the nfExM protocol—gelation, softening, immunostaining, and expansion in water. Immunostaining for microtubules was performed using rabbit anti-β-tubulin primary antibody (Abcam, Catalog no. ab6046; dilution: 1:100), followed by incubation with goat anti-rabbit secondary antibody conjugated to Alexa Fluor 546 (Thermo Fisher, Catalog no. A11035; dilution: 1:100). After expansion, microtubule structures were imaged with a spinning disk confocal microscope (see **Image Acquisition** section for microscope details). Structural continuity and integrity of microtubules were qualitatively assessed by visually examining acquired images for the presence or absence of discontinuities along microtubule filaments.

### UV Photoinitiation and Measurement of UV Power Density at the Sample Plane

Given the failure of APS/TEMED to initiate polymerization at –20 °C, we adopted a UV-induced polymerization strategy using Irgacure 2959 (I-2959; Sigma-Aldrich, Catalog no. 410896), a temperature-independent photoinitiator previously demonstrated to rapidly induce gelation in acrylamide/sodium acrylate-based systems at room temperature^23^. We prepared a monomer solution consisting of 46.7% w/w N,N-dimethylacrylamide (DMAA), 9.7% w/w sodium acrylate (SA), and 0.1% w/w N,N’-methylene-bis-acrylamide (Bis) in ultrapure water, supplemented with 4.9% w/w I-2959. Polymerization was initiated by UV illumination. For standard experiments, we used a handheld UV lamp (Analytik Jena, Catalog no. 95-0343-01) emitting at a central wavelength of 365 nm. The UV lamp was positioned approximately 4 cm above the sample surface.

The UV power density at the sample plane was measured as 0.0171 mW/mm² using a power meter (Thorlabs S120C sensor and Thorlabs PM100A console), placed directly beneath a No. 2 cover glass (VWR, Catalog no. 48382-136) to replicate experimental conditions accounting for optical attenuation by the glass. We also evaluated polymerization efficiency using an alternative handheld UV source (Grafco E-Series, Model 2214), positioned 1 cm from the sample. Under these conditions, the UV power density was measured aa 0.0348 mW/mm², again using the Thorlabs power meter with the No. 2 cover glass positioned between the lamp and the sensor. Polymerization was successfully achieved within 1 minute under these conditions. However, for consistency with previously acquired data, subsequent experiments exclusively utilized the Analytik Jena lamp configuration with the 0.0171 mW/mm² power density.

### Fixation, gelation, and denaturation of cell culture with proExM(4% PFA fix)

Reagents being used are 16% paraformaldehyde (PFA; EMS, Catalog no. 15710), Glycine (Sigma Aldrich, Catalog no. G7126-100G), Acryloyl-X, SE (AcX; Thermo Fisher, Catalog no. A20770), Sodium acrylate (Pfaltz & Bauer, Catalog no. S03880), Acrylamide (Sigma Aldrich, A9099-25g), N,N′-methylenebisacrylamide (Sigma Aldrich, 146072), Sodium chloride (NaCl; Thermo Fisher, AM9759), Ammonium persulfate (APS; Thermo Fisher, Catalog no. 17874), N,N,N′,N′-Tetramethylethylenediamine (TEMED; Thermo Fisher, Catalog no. 17919), 20% Sodium n-dodecyl sulfate (SDS; Thermo Fisher, AM9820), Tris(hydroxymethyl)aminomethane (Tris; Thermo Fisher, AM9855G).

Cells were rinsed once at RT with Dulbecco’s Phosphate-Buffered Saline (DPBS, Thermo Fisher, Catalog no. 14190136) prewarmed to 37 °C. Then cells were fixed with freshly prepared 4% w/v paraformaldehyde (PFA; EMS, Catalog no. **1.** 15710) in PBS for 10 min at room temperature (RT). After fixation, the fixative was carefully removed, and residual PFA was quenched by incubation in 100 mM glycine (Sigma Aldrich, Catalog no. G7126-100G) in PBS for 10 min at RT. Cells were then washed three times with PBS for 5 min each. Fixed cells were anchored by incubation in 0.1 mg/mL AcX (Acryloyl-X, SE; Thermo Fisher, A20770) in PBS at RT for 2 hours. After anchoring, the AcX-treated cells were washed twice with PBS for 5 min each at RT. The monomer solution for gelation was prepared with 8.625% w/v sodium acrylate, 2.5% w/v acrylamide, 0.15% w/v N,N′-methylenebisacrylamide, and 2 M NaCl in PBS. This monomer solution was prepared in advance, frozen in aliquots, and thawed immediately prior to use. To initiate polymerization, APS and TEMED were added to the monomer solution as initiator and accelerator, respectively, at a final concentration of 0.15% w/v each with APS added last. The monomer solution was added to the cells and gelation was carried out in a humidified incubator at 37°C for 1.5 hours. After polymerization, the gel was carefully transferred to denaturation buffer containing 200 mM SDS, 200 mM NaCl, and 50 mM Tris (pH 9.0). Denaturation was performed at 95°C for 1.5 hours. Following denaturation, the gels were washed thoroughly with PBS to remove residual SDS. The expanded gels were incubated with primary antibodies against β-tubulin (ab6046, with 1:100 dilution) overnight at 4°C. After washing three times with PBS, 30 min each at RT, samples were incubated with goat anti-rabbit secondary antibody conjugated to Alexa Fluor 546 (Thermo Fisher, Catalog no. A11035; dilution: 1:100) for 5 hours at RT. Samples were washed again three times with PBS, 30 min each at RT. The immunostained gels were expanded in excess distilled water until fully expanded.

### Fixation, gelation, and denaturation of cell culture with nfExM

Cells were rinsed once at RT with DPBS prewarmed to 37 °C, then immediately supplied with acrolein-methanol solution (Restek, Catalog no. 30645), previously chilled to −20°C, and incubated for 8 min at −20°C (see **Supp. Note 3** for details on performing this and following nfExM steps while maintaining the sample at – 20 °C). Subsequently, nfExM monomer solution, previously chilled to −20°C, was added to the cells without any intermediate washing step. The cells were then incubated for 5 minutes at −20°C, allowing sufficient time for the monomer solution to permeate the cellular structure. Then approximately 27 μL of the monomer solution (−20°C) was added onto the slide of a gelation chamber, which was prepared with parafilm strips around a slide pre-chilled at −20°C (see **Supp. Fig. 3** for chamber construction). A syringe needle with a bent tip along with forceps was used to carefully lift and transfer the coverslip with cells from a 24-well plate to the gelation chamber, and the cell-coverslip was placed facing down to close the chamber. Finally, gelation was induced using a UV light (with 0.0171 mW/mm² power density) with an exposure time of 3 minutes at −20°C .

After gelation, gels were incubated in MAP denaturation buffer (200 mM SDS, 200 mM NaCl, and 50 mM Tris, with the final pH adjusted to 9.0) for 1.5 hours at 95°C followed by washing three times with 1x PBS for 30 minutes each to remove any residual denaturation buffer and subject to immunostaining.

### Fixation, gelation, and denaturation of cell culture with CryoExM

Reagents being used were paraformaldehyde (PFA; EMS, Catalog no. 15710), acetone (Thermofisher Scientific, Catalog no. AC326801000), Acrylamide (Sigma Aldrich, Catalog no. A9099-25G), Bis-acrylamide (Sigma-Aldrich, Catalog no. 146072), SA (Pfaltz & Bauer, Catalog no. S03880), ammonium persulfate (APS, Thermo Fisher, Catalog no. 17874), N,N,N′,N′-Tetramethylethylenediamine (TEMED; Thermo Fisher, Catalog no. 17919), nuclease-free water (Thermo Fisher Scientific, Catalog no. 10977015)

The plunge-freezing and freeze substitution were performed as previously described^19^. To enhance the consistency of our results, we opted for a commercially manufactured cryoplunger, the Gatan CP3, instead of the homemade plunger. The vitrified cells were prepared by positioning a No. 1.5 12-mm coverslips, bearing the biological specimen, using precision tweezers to hold them at their midpoint. Excess culture medium was carefully blotted away with filter paper before the coverslips were submerged into liquid ethane using the Gatan CP3 cryoplunger. Subsequent to plunge-freezing, the coverslips were immediately transferred into a pre-chilled 5-ml Eppendorf tube filled with solidified acetone (Thermofisher Scientific, Catalog no. AC326801000) at liquid nitrogen temperature. To ensure optimal cryofixation, the tubes were positioned at a 45-degree angle on dry ice and gently agitated overnight, allowing a gradual temperature increase to 0°C. Rehydration was performed at room temperature in a graded ethanol–water series, 5 min in each incubation: 100% ethanol (twice; stock stored at −20 °C), 95% v/v ethanol (twice; stock stored at −20 °C), 70% v/v ethanol (stock stored at 4 °C), 50% v/v ethanol (stock stored at 4 °C), and finally PBS at room temperature. The cells were then preserved in PBS for further expansion microscopy process.

The cryofixed cells were incubated in a solution of 2% AA and 1.4% w/v formaldehyde in PBS for a duration of 4 hours at a temperature of 37°C. This step was followed by incubation in a CryoExM monomer solution (19% w/w SA, 10% w/w AA, and 0.1% w/w Bis in PBS) for 5 minutes on ice, just after the addition of 0.5% w/v TEMED and 0.5% w/v APS to the solution. Gelation was performed in a humidified 37 °C incubator for 1 h. After gelation, gels were incubated in MAP denaturation buffer (200 mM SDS, 200 mM NaCl, and 50 mM Tris, with the final pH adjusted to 9.0) for 1.5 hours at 95°C followed by washing three times with 1x PBS for 30 minutes each at RT to remove any residual denaturation buffer. Sample were stored at 4°C or subjected to immunostaining in the next step.

### Fixation, gelation, and denaturation of cell culture with high T% proExM

Cells were fixed with freshly prepared 4% w/v PFA in PBS for 10 min at RT. After fixation, the fixative was carefully removed, then incubate in 100 mM Glycine in PBS for 10 min at RT. Next, cells were washed thoroughly three times with PBS for 5 min each at RT. Fixed cells were anchored by incubation in 0.1 mg/mL AcX in PBS for 2 hours at RT. After anchoring, the AcX-treated cells were washed twice with PBS for 5 min each at RT. The nfExM monomer solution, pre-chilled to −20°C, was added to the cells and incubated for 5 minutes at −20C. The cells were then transferred to a gelation chamber (prepared as described above) and gelation was initiated using the UV light (with 0.0171 mW/mm² power density) for 3 minutes at −20C. After gelation, gels were incubated in MAP denaturation buffer (200 mM SDS, 200 mM NaCl, and 50 mM Tris, pH 9.0) for 1.5 hours at 95°C followed by washing three times with PBS for 30 minutes each to remove residual denaturation buffer. Post-expansion immunostaining and imaging procedures followed the same protocol as described for nfExM treatment.

### Cold PFA Fixation, gelation, and denaturation of cell culture

Cold fixation solution was prepared one day before use by diluting 32% w/v PFA (EMS, Catalog no. 15714) aqueous solution with anhydrous methanol to a final concentration of 4% w/v and the solution stored at −20°C. Cells were fixed with cold 4% w/v PFA in methanol for 10 minutes at −20°C. After fixation, the fixative was carefully removed and cells were washed thoroughly three times with PBS for 5 min each at RT. Fixed cells were anchored by incubation in 0.1 mg/mL AcX in PBS at RT for 2 hours. After anchoring, the AcX-treated cells were washed twice with PBS for 5 min each at RT. Subsequently, cells were processed using either the standard proExM protocol or the high T% proExM protocol as described above.

### Fixation, gelation, and denaturation of cell culture with proExM/dExPath/MAP style ExM

Reagents being used are glutaraldehyde (EMS, Catalog no. 16019), paraformaldehyde (PFA; EMS, Catalog no. 15710), Acrylamide (Sigma Aldrich, Catalog no. A9099-25G), Bis-acrylamide (Sigma-Aldrich, Catalog no. 146072), SA (Pfaltz & Bauer, Catalog no. S03880), ammonium persulfate (APS, Thermo Fisher, Catalog no. 17874), N,N,N′,N′-Tetramethylethylenediamine (TEMED; Thermo Fisher, Catalog no. 17919), nuclease-free water (Thermo Fisher Scientific, Catalog no. 10977015), sodium borohydride (Sigma Aldrich, Catalog no. 213462-25G), Glycine (Sigma Aldrich, Catalog no. G7126-100G), Tris(hydroxymethyl)aminomethane (Tris; Thermo Fisher, Catalog no. AM9855G), Sodium chloride (NaCl, Thermo Fisher, Catalog no. AM9759), Sodium hydroxide(NaOH; Sigma, Catalog no. S8263), Sodium n-dodecyl sulfate (SDS; Thermifisher, Catalog no. AM9820), EDTA (ThermoFisher, Catalog no. AM9260G), beta mercaptoethanol(ThermoFisher, Catalog no. 35602BID).

For the fixation of microtubules, cells were rinsed once with PEM buffer (0.1 M PIPES, 1 mM EGTA, 1 mM MgCl2, 0.4% w/v Triton X-100, pH 7.0) for 45 s at RT, and then fixed with freshly prepared fixation solution (3% w/v PFA/0.1% w/v GA in PBS) for 10 min at RT. Then fixatives were removed, and the cells were incubated in freshly prepared 0.1% w/v sodium borohydride in PBS for 7 min at RT. The cells were incubated with 100 mM glycine in PBS for 10 min at RT, then washed three times with PBS for 5 min each at RT.

After fixation, cells were incubated in 0.1 mg/mL AcX in PBS for 4 hours at RT. AcX-treated cells were washed twice with PBS for 5 min each at RT, and the cells were then expanded via the following steps (gelation, digestion and expansion). Monomer solution (2 M NaCl, 8.625% w/v SA, 2.5% w/v AA, 0.15% w/v Bis, in PBS) was mixed, frozen in aliquots, and thawed before use. The initiator APS and the accelerator TEMED were added to the monomer solution to 0.2% w/v each with APS added last. Gelation was performed in a humidified 37 °C incubator for 1.5 h.

After gelation, gels were underwent different denaturation treatments based on the protocol under test: proExM: 100 mM Tris base, 5% w/v Triton X-100, 1% w/v SDS autoclaved at 120 °C for 1 h; dExPath: 20% w/v SDS, 50 mM Tris, 25 mM EDTA, 0.5% w/v Triton X-100, 100 mM beta mercaptoethanol, autoclaved at 120 °C for 1 h; MAP: 200 mM SDS, 200 mM NaCl, and 50 mM Tris in DI water (pH adjusted to 9.0) at 95 °C for 1.5 h. Then the gels were thoroughly washed three times with PBS for 30 min each and subject to immunostaining.

### Post-expansion immunostaining of expanded gels

Followed by denaturation, the gels were incuabted in MAXblock blocking medium (Active Motif, Catalog no. 15252) for 30 minutes to 1 hour at RT. After blocking, the gels were washed once with MAXwash washing solution (Active Motif, Catalog no. 15254) at RT for 10min, followed by staining with 1:100 (for Fig. 2 and Fig. 5) or 1:200 (to economize reagent, for Fig.3 and Fig. 4) diluted primary antibodies (for tubulins, Abcam, ab6064; for TOM20, Abcam, ab186735; for GM130 Proteintech, 11308-1-AP) overnight at 4 °C. After primary antibody staining, the gels were washed four times with MAXwash for 30min each at RT, followed by staining with 1:100 (for Fig. 2 and Fig.5) or 1:200 (for Fig.3 and Fig. 4) diluted secondary antibodies (for tubulins: Alexa Fluor 546, Thermo Fisher, Catalog no. A11035; for TOM20 and GM130: Alexa Fluor 488 Thermo Fisher, Catalog no. A11008) for 5h at RT. Both primary and secondary antibodies were diluted in MAXbind staining medium (Active Motif, Catalog no. 15253). For samples stained with GM130, gels were additionally stained with Hoechst 33342 (Thermo Fisher, Cat. No. 62249) following secondary antibody incubation, by incubating in 4 µM Hoechst in PBS for 10 min at RT, to provide a nuclear counterstain for cell identification. The gels were then washed three times with MAXwash for 30min each at RT, followed by washing with excess distilled water at RT until fully expanded.

### Expansion Factor Measurement

After gelation, each gel was manually trimmed with a razor blade into an approximately 5 mm × 5 mm square shape, with one corner diagonally removed (shortened to ∼4 mm, **Fig. 1fii**) to create an asymmetric fiducial marker for orienting the cell-containing region during imaging. Following denaturation, immunostaining, and full expansion in excess deionized water, the gel width was measured using a ruler. The linear expansion factor was calculated as the post-expansion dimension divided by the original trimmed width of 5 mm, measured from four independent gels from one batch experiment. This expansion factor was used to convert expanded measurements back to biological units for downstream quantification.

### Image Acquisition

All fluorescence imaging for Figures 2–5 was performed using a CSU-W1 spinning-disk confocal microscope (Nikon) equipped with a SoRa module, operated in standard confocal mode; SoRa super-resolution mode was not used. Images for Supp. Tables 1 and 2 were acquired on a separate Nikon CSU-W1 spinning-disk confocal system. Images were acquired with a Nikon CFI Apochromat LWD Lambda S 40XC WI water immersion objective lens (40x, numerical aperture 1.15, working distance 0.59–0.61 mm) unless otherwise specified.

Fluorescence imaging was performed using three excitation wavelengths: 405 nm (Hoechst nuclear counterstain in Figure 5, co-stained with GM130), 488 nm (Alexa Fluor 488 in Figure 5 and Supp. Figure 1), and 561 nm (Alexa Fluor 546 in Figures 2–4). Emission was collected through the following bandpass filters: 455/50 nm (405 nm channel), 525/36 nm (488 nm channel), and 605/52 nm (561 nm channel).

To ensure quantitative comparability within each experiment, all images grouped within individual figures (except Supp. Tables 1 and 2, which show qualitative examples for protease and cleavable crosslinker screening rather than quantitative comparisons) were acquired using the same imaging parameters: laser power, exposure time, objective lens, excitation wavelengths, and emission filters. For display purposes, brightness and contrast were adjusted using Fiji’s automated adjustment function followed by manual refinement (raising minimum and lowering maximum intensity thresholds) to enhance visualization of features of interest. Adjustments were linear and applied uniformly to prevent bias. For Figure 2 main panels and their corresponding insets, the minimum and maximum display intensity thresholds were set the same within each image type (main panels or insets), though the ranges differed between main panels and insets. All quantitative analyses of fluorescence intensities were performed on unprocessed raw image data, as specified in Results and figure legends.

### Quantitative Analysis of Microtubule Structural Continuity with Local Adaptive Threshold Method

Microtubule gap analysis was performed using a custom sliding-window algorithm to detect local intensity discontinuities. Individual microtubules were manually traced using Fiji ImageJ free hand tool, with microtubule segments that were not overlapped with others to avoid intensity increases due to overlapping segments. In Fig. 4, microtubule segments were traced only in regions without visible gel cracks. The line tool was configured with a 3-pixel perpendicular sampling width, which accounts for the width of microtubules while minimizing inclusion of surrounding background signal. Intensity profiles were extracted for each traced microtubule segment using the “Plot Profile” function in Fiji, which provided intensity values at each pixel position along the traced line. This approach generates intensity profiles where each data point represents the mean fluorescence intensity across the microtubule cross-section, reducing the impact of random fluctuations in signal intensity that might occur with single-pixel line width measurements.

Background signal measurements were obtained from areas devoid of microtubule structures, and for each image, both the mean background intensity and standard deviation were calculated to ensure accurate baseline correction. Image-specific background correction was applied using the equation I_corrected[i] = I_raw[i] - μ_bg, where I_raw[i] represents the fluorescence intensity value at position i along the traced microtubule, and μ_bg represents the background mean measured from regions devoid of microtubule signal. The algorithm employed a locally adaptive threshold method with a symmetric sliding window of ±3 pixels around each position i. Window boundaries were dynamically adjusted for edge positions using: Window_start = max(1, i-3) and Window_end = min(n, i+3), where n represents the total number of data points along the traced segment.

For each position, the local mean intensity μ_local[i] was calculated within the sliding window, and position-specific thresholds were determined as Threshold[i] = μ_local[i] × α, where α represents the relative intensity threshold parameter. Positions were classified as gaps when I_corrected[i] < Threshold[i]. Primary analysis employed α = 0.8, with threshold sensitivity analysis performed by varying α from 0.6 to 0.9 and quantifying gap ratio and size at each value to evaluate the effect of parameter choice on measured gap metrics.

Each data point along the intensity profile was then binary-coded, where points with intensity values below the threshold were designated as ’1’ (gap), and points with intensity values above the threshold were designated as ’0’.

The gap ratio for each microtubule segment was calculated by dividing the total number of points identified as gaps (sum of ’1’s) by the total number of data points (sum of ’1’s and ’0’s) along the segment: Gap Ratio = (Total number of ’1’s) / (Total number of data points). For analyzing the physical dimensions of gaps, consecutive ’1’s in the binary-coded intensity profile were grouped together into single gap segments. The size of each gap segment was calculated by multiplying the number of consecutive ’1’s (representing a single gap segment) by the pixel size at the post-expansion scale, and then dividing by the expansion factor to convert to biological units.

### Quantitative analysis of mitochondrial and Golgi structural preservation

Expanded samples were imaged by spinning disk confocal microscopy. Samples were immunostained for GM130 (cis-Golgi marker) or TOM20 (outer mitochondrial membrane protein) and labeled with Alexa Fluor 488–conjugated secondary antibodies. GM130-stained samples were additionally counterstained with Hoechst to label nuclei, whereas TOM20-stained samples were imaged without nuclear staining. Z-stacks were acquired for each field of view and processed using custom Python scripts based on the nd2 library. For GM130 datasets, the 488 nm channel was extracted and converted into a two-dimensional maximum-intensity projection (MIP) over all acquired z-planes; TOM20 datasets were likewise projected along the z-axis. All subsequent analyses were performed on these MIP images.

Automated image segmentation and quantification were performed to assess structural preservation.

1. **Background Correction.** For each image, background fluorescence was estimated as the 10th percentile of the raw pixel intensities and subtracted to generate background-corrected images.
2. **Structure Mask Generation.**

### 2.1. Binary segmentation

Structural regions of interest (ROIs) were defined by applying a fixed intensity threshold of 50 arbitrary units (a.u.) to the corrected images to create initial binary masks. This threshold value was selected based on preliminary visual inspection to achieve robust segmentation of visible structures.

To validate this choice and assess potential threshold-dependent biases, we performed a sensitivity analysis testing thresholds of 30, 50, 100, and 150 a.u. on the background-corrected images (Fig. 5i–l). While the absolute signal pixel counts and mean intensities changed as expected with threshold variation— higher thresholds exclude dimmer pixels, resulting in lower pixel counts but higher mean intensities—the relative differences between nfExM and CryoExM remained consistent. Across all tested thresholds, nfExM exhibited higher signal retention and intensity compared to CryoExM for both TOM20 and GM130. This confirms that our conclusions regarding nfExM’s superior protein retention are robust to threshold selection and not dependent on a specific parameter value.

### 2.2. Noise removal size cutoff determination

The initial binary masks were then refined using scikit-image (v0.21+) connected-component analysis to remove small objects below size thresholds of 100 pixels for Golgi and 120 pixels for mitochondria. These size cutoffs were determined based on the known minimum dimensions of these organelles. For mitochondria, the smallest reported diameter is approximately 0.5 µm^39^ in biological samples, corresponding to a cross-sectional area of ∼0.196 µm². After 4-fold linear expansion, this corresponds to ∼3.14 µm² in expanded space. At our imaging resolution of 0.1625 µm/pixel, this translates to approximately 119 pixels, justifying our 120-pixel cutoff.

For Golgi, the diameter of the cisternae typically range from 0.7–1.1 µm in diameter^40^, with individual cisternae often appearing as smaller punctate or fragmented structures after immunostaining and expansion (Fig. 5c, d). To avoid excluding genuine but small Golgi elements—particularly in CryoExM samples where fragmentation was more apparent—we used a more conservative (lower) cutoff of 100 pixels compared to the calculated 233 pixels. This conservative approach ensures retention of dim or fragmented genuine Golgi structures while still excluding obvious imaging artifacts such as single-pixel noise or small non-specific fluorescent debris.

### 2.3. Hole filling

Interior gaps within structures were filled using scikit-image’s binary_fill_holes function to generate continuous masks. This step corrects for local intensity heterogeneities within organelles that can arise from antibody penetration variability.

## 3. Quantification metrics

Structural preservation was quantified using two metrics. The signal pixel count was defined as the total number of mask pixels with positive intensity after background subtraction. The mean fluorescence intensity was defined as the average background-corrected intensity across all pixels within each structure mask. Data processing and statistical analysis were conducted using NumPy (v1.24+), pandas (v2.0+), and SciPy, with visualizations generated using matplotlib (v3.7+).

## Code availability

The custom Python scripts used for image processing and quantification are available at https://github.com/yixiliumit/nfExM-analysis. The code is released under the MIT license.

## Contributions

Y.L. and C.Z. initiated exploratory studies of cleavable fixatives for ExM and contributed ideas and experiments during the early phase of the project. Y.L. subsequently conceived and developed nfExM, designed and performed the experiments for its development and validation, and analysed and interpreted the resulting data. Y.L. and C.Z. analysed and interpreted data from the early cleavable-fixative studies. Y.L., C.Z. and C.S. wrote and edited the manuscript. E.S.B. supervised the overall project, initiated the work, contributed key ideas, designed experiments, helped with data analysis and interpretation and edited the manuscript.

## Acknowledgments

ESB acknowledges support by Ashar Aziz, Lisa Yang and Hock Tan, the Alana Center at MIT, NIH R01GM160726, W.M. Keck Foundation, Richard King Mellon Foundation, NIH R01AG087374, NIH 1R01EB024261, Tom Stocky and Avni Shah, Kathleen Octavio, Lore McGovern, Jed McCaleb, James Fickel, Good Ventures/Open Philanthropy, NIH 1R01AG070831, the European Research Council (ERC) under the European Union’s Horizon 2020 research and innovation programme (grant agreement No 835102), John Doerr, and HHMI; and in part by the Koch Institute Support (core) Grant P30-CA014051 from the National Cancer Institute. Y.L. acknowledges support from the Alana Fellowship during doctoral work contributing to this study and from the J. Douglas Tan Postdoctoral Fellowship. We thank D. S. Yun and the Nanotechnology for Integrative BioEM Core at the Koch Institute’s Robert A. Swanson (1969) Biotechnology Center for assistance with plunge freezing and technical support. We thank D. Estandian, A. Choueiri, A. Wassie, T. Shin and M. Karagiannis for helpful discussions.

