## Supplementary Information for "Network fixation expansion microscopy"

### **Supplementary Notes**

#### **Supplementary Note 1: Acrolein Handling and Quality**

The acrolein-methanol solution (Restek, Cat. No. 30645) has a manufacturer-stated shelf life of 3 months. We had experience where solutions used one month past the expiration date produced poor microtubule preservation quality, similar to that observed with PFA fixation (Fig. 3a). However, we did not systematically characterize how preservation quality degrades as a function of time relative to the expiration date. We therefore recommend to always use fresh solutions that are well within their shelf life.

The manufacturer provides the acrolein-methanol solution in a glass ampule containing 1 mL of solution. This relatively small volume, even if pre-chilled to $-20^{\circ}\text{C}$, can be rapidly warmed well above $-20^{\circ}\text{C}$ simply by holding the ampule in one's hand. Therefore, prior to each experiment, we transferred the required volume of acrolein-methanol solution from the original glass ampule into an amber microcentrifuge tube in the fume hood, and then chilled the tube to $-20^{\circ}\text{C}$ in a metal cold block before use. When retrieving the solution for use, the amber tube was kept fully seated in the cold block at all times, with only the tube cap being opened (but not lifting the tube from the block) to withdraw the liquid. After adding the acrolein solution to the 24-well plate, we sealed the well with laboratory tape or adhesive plate sealing film to prevent acrolein evaporation during incubation.

#### **Supplementary Note 2: Sub-pixel gap detection limits**

Our algorithm works as follows: for each pixel, we take the average intensity of a 7-pixel window (the pixel itself plus 3 on each side), and mark it as a gap if the pixel is at least 20% dimmer than that average (i.e., its intensity $< 0.8 \times$ local mean).

The smallest gap we can report is 1 pixel ($0.041\ \mu\text{m}$). What happens to a gap smaller than 1 pixel depends on where it lands on the pixel grid — see **Supp. Figure 4** for three scenarios (using a 0.3-pixel-wide gap as an example for scenarios B and C). For illustration we assume a uniform intensity of 100 along an intact microtubule; real data has fluctuations from noise and staining variation, but the alignment-dependent behavior we describe holds regardless.

• **Scenario A:** A gap that fills roughly one whole pixel drops that pixel's intensity close to zero, which easily crosses the threshold. Reported as a 1-pixel gap.
• **Scenario B:** A sub-pixel gap (e.g., 0.3 pixel wide) that sits inside one pixel still drops that pixel's intensity enough (e.g., from 100 to 70) to cross the threshold. Also reported as a 1-pixel gap — even though the real gap is smaller.
• **Scenario C:** The same 0.3 pixel wide gap that spanning two pixels splits the intensity drop between them. Neither pixel drops enough to cross the threshold, so the gap is missed entirely (reported as 0).

So a 1-pixel gap in our data could be a real 1-pixel gap (scenario A), or a sub-pixel gap that happened to land inside one pixel (scenario B). Gaps that crossed pixel boundaries (scenario C) don't show up at all, if too small.

#### **Supplementary Note 3: Performing fixation and gelation at –20 °C**

To ensure the sample and reagents were maintained at –20 °C for the duration of the nfExM fixation, monomer incubation, and gelation steps (see Methods section, “Fixation, gelation, and denaturation of cell culture with nfExM,” for details of the nfExM treatments), we typically did the following:

• Beginning just after the addition of –20 °C acrolein-methanol solution, the 24-well plate containing coverslip with cells was placed and kept on a pre-chilled thermal workstation with a metal surface (Corning CoolBox XT 432021 with Corning XT Freezing Core 432082 and Corning CoolSink XT96F 432070) to hold the plate and sample at –20 °C as it was moved between steps.
• To perform the –20 °C incubations (in acrolein-methanol or monomer solution), the entire well plate-thermal workstation assembly was moved into a –20 °C freezer.
• Before adding monomer solution and coverslip with cells, the gelation chamber slide was pre-chilled and maintained at –20 °C by placing it on a separate metal cold block pre-chilled to –20 °C.
• The 27 µL drop of monomer solution was placed on this chilled gelation chamber immediately (5–20 s) prior to closing the chamber with the coverslip containing cells.
• The closed gelation chamber along with the metal cold block was immediately (within 60 s) transferred to a –20 °C freezer.
• The UV exposure was then performed by placing the UV exposure unit (**Supp. Fig. 3**) directly above the gelation chamber inside of the –20 °C freezer.

Alternatively, this entire procedure can be carried out in a –20 °C cold room if one is available.

#### Supplementary Figures

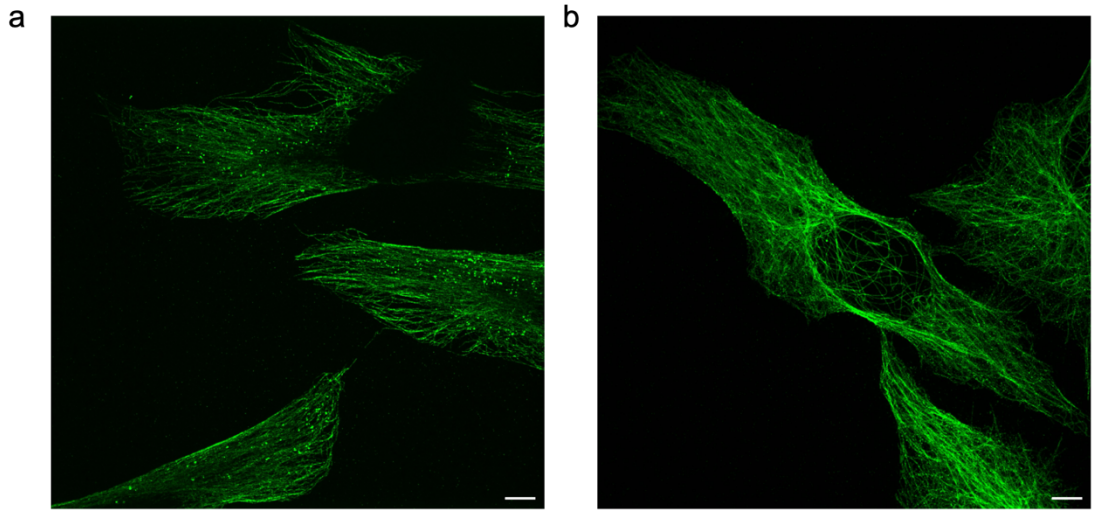

**Supplementary Figure 1. Cold processing steps alone (PFA at -20 °C) are insufficient for microtubule preservation, even with high gel monomer concentration.**

To test whether nfExM's improvement in protein retention comes from performing steps at low temperature rather than from its acrolein chemistry, we added cold PFA fixation steps to the proExM protocol (which uses Acryloyl-X, SE (AcX) anchoring), without introducing acrolein chemistry. These were initial tests using a single culture batch; observations are thus preliminary, but helped guide us along the path.

(a) Representative ($n = 6$ images from 1 culture batch) confocal image (maximum intensity z-projection) of expanded HeLa cells, post-expansion immunostained for β-tubulin. Protocol: 4% w/v paraformaldehyde (PFA) in methanol at -20°C for 10 min, followed by AcX anchoring at room temperature for 2 hrs, followed by gelation of a standard proExM gel (low total monomer concentration, or low T%, where T% denotes the total monomer percentage in units of w/v, in this case 11.1% w/v) at 37°C for 1.5 h.

(b) Representative ($n = 9$ images from 1 culture batch) confocal image, as in (a), but with fixation with 4% w/v PFA in methanol at -20°C for 10 min, followed by AcX anchoring at room temperature for 2 hrs, followed by gelation of an nfExM high T% (59.2% w/v) monomer solution at -20°C via UV initiation for 3 min.

All images were acquired under the same microscope settings with a 40x 1.15 NA water objective (Nikon). Images were visualized with the auto-scaling function in Fiji. Scale bar (in biological units, e.g. real space units divided by expansion factor): 5 μm.

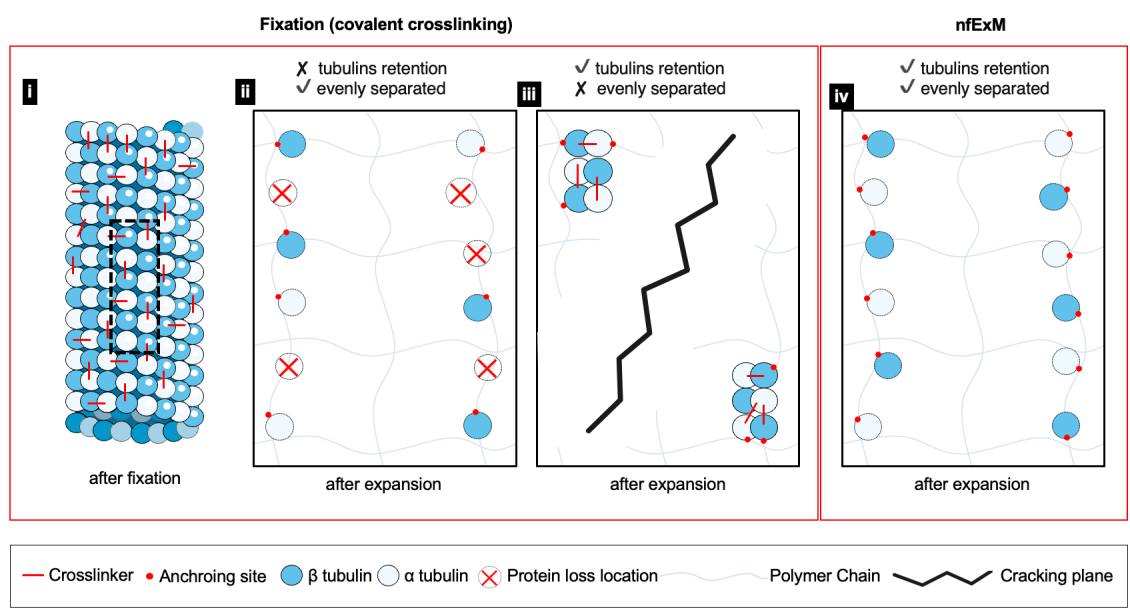

### Supplementary Figure 2. Proposed schematic model for how fixation strategy determines microtubule structural outcomes in ExM.

This schematic proposes that the strength of fixation-induced protein-protein crosslinks determines how microtubule organization is preserved after expansion. Our proposed model can be used to interpret the results of three scenarios: weak fixation (panels i and ii, e.g., PFA), strong fixation (panels i and iii, e.g., GA), and protein-network fixation (panel iv, nfExM).

i) Schematic of microtubule organization after standard covalent crosslinking. Tubulin heterodimers (blue circles, β-tubulin; white circles, α-tubulin) are crosslinked (red lines) to neighboring subunits to preserve nanoscale organization. The strength of these protein-protein crosslinks determines subsequent expansion outcomes. Black dashed line box shows the region of interest highlighted in panels ii, iii, and iv.

(ii) Weak fixation scenario (e.g., PFA): expansion with weak protein-protein crosslinking leads to the loss of many tubulin subunits (red X, protein loss locations), although the gel expands uniformly (✓ evenly separated). This is consistent with the reduced intensity and increased gap size and ratio observed in standard proExM and CryoExM (Fig. 2d–f).

(iii) Strong fixation scenario (e.g., GA): expansion with extensive protein-protein crosslinking. Glutaraldehyde creates rigid protein assemblies (tubulins at the top left and bottom right) that resist protein loss (✓ tubulin retention) but impede uniform gel swelling. Mechanical stress from these rigid assemblies could generate visible cracks (zigzag, cracking plane) and non-uniform expansion (✗ evenly separated; deformed polymer chains) during expansion. This is consistent with good microtubule continuity (Fig 4o–p) but frequent cracking (Fig 4t) in GA-based methods.

(iv) nfExM strategy: Tubulin subunits are anchored directly to the polymer network.

During expansion, proteins move with the gel, achieving both high retention (✓ tubulin retention) and uniform separation (✓ evenly separated). This is consistent with nfExM's minimal gaps (Fig 4o–p), yet zero cracking (Fig 4t).

This schematic offers a hypothesis for how fixation strategy shapes ExM outcomes: weak fixation preserves gel uniformity but loses protein; strong conventional crosslinking preserves protein but causes cracking; our nfExM strategy (anchoring proteins directly to the gel) retains both protein and gel uniformity. The model is consistent with our observed intensity, gap, and cracking measurements (Figs. 2, 3, 4), but direct biophysical validation (e.g., gel mechanics, protein-gel binding forces) would be needed to test these mechanisms.

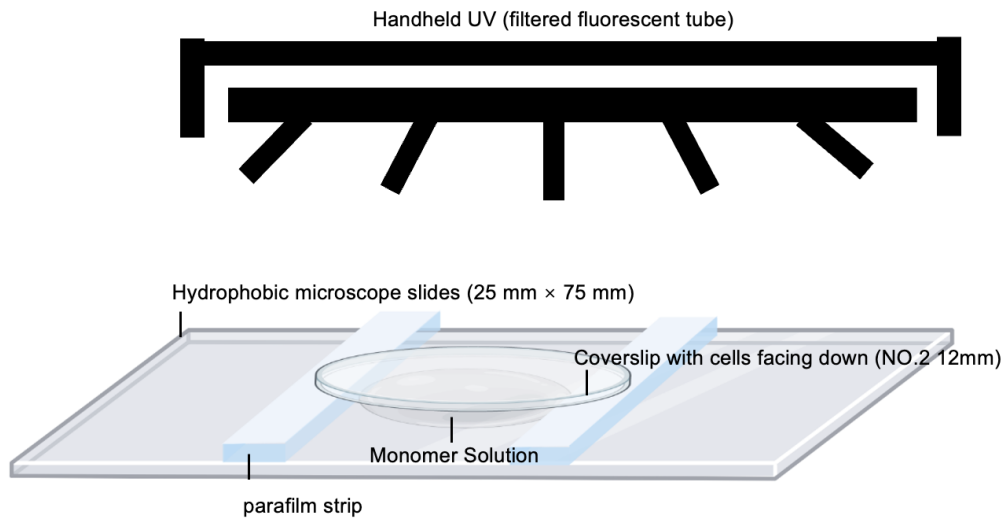

#### **Supplementary Figure 3. Schematic Illustration of gelation chamber assembly and UV-initiated polymerization setup for nfExM.**

The schematic illustration depicts the components of the custom-built gelation chamber used for nfExM. The chamber consists of hydrophobic microscope slides (25 mm × 75 mm) serving as the base, which provides a non-adherent surface for the polymerized gel to facilitate peeling the polymerized gel from the slide in a later step; see Methods section, “Gelation Chamber Preparation” for the silane-based hydrophobic treatment protocol. Note: as a simpler alternative, wrapping the slide surface with a layer of Parafilm prior to chamber assembly also provides a sufficient non-adherent surface for gel release without chemical treatment. A monomer solution is placed at the center of two spacer strips ( ~ 27 μL for cells cultured on 12 mm diameter coverslips), and a No. 2 coverslip containing the cells is positioned cell-side down on top of the monomer solution to close the chamber. To control the gel thickness, the coverslip is supported by two parallel Parafilm strips (or a No.2 coverslip) placed ~10 mm apart (for a 12 mm diameter coverslip). This spacing defines the final thickness of the polymerized gel. The chamber is then exposed to UV light emitted from a handheld UV lamp (Analytik Jena, catalog number #95-0343-01) with a center wavelength of 365 nm, positioned approximately 4 cm from the sample. The UV power density at the sample was measured at 0.0171 mW/mm<sup>2</sup> using a power meter (Thorlabs), with an exposure duration of 3 minutes. The entire assembly is maintained at -20°C until the polymerization process is complete, typically by working inside a -20°C freezer or a -20°C cold room.

**A. Real gap ≈ 1 pixel (fully occupies one pixel)**

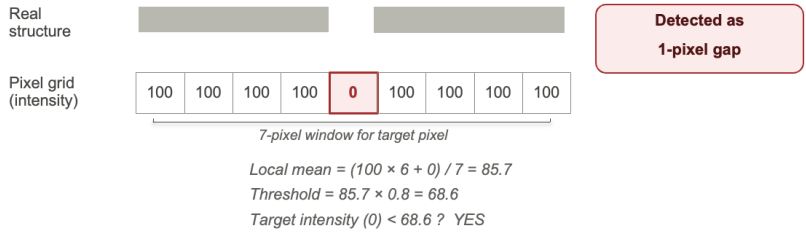

**B. Real gap = 0.3 pixel, sitting inside a pixel**

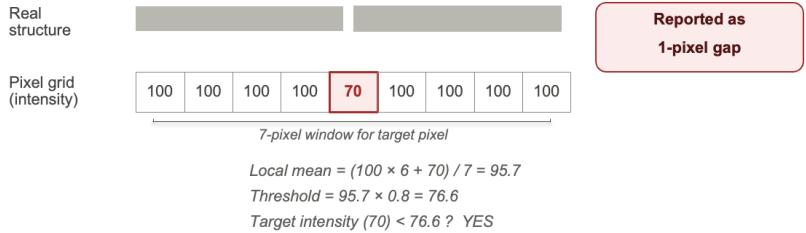

**C. Real gap = 0.3 pixel, crossing a pixel boundary**

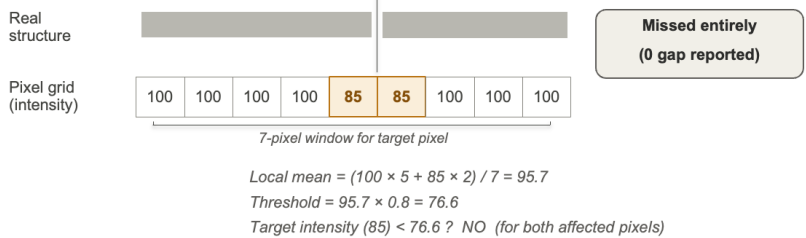

**Supplementary Figure 4. How sub-pixel gaps appear in pixel-by-pixel detection.**

Three scenarios showing why a gap smaller than 1 pixel can either show up as a 1-pixel gap or be missed entirely, depending on how it aligns with the pixel grid. Scenarios B and C use a 0.3-pixel-wide gap as an example. For illustration, gap-free microtubule regions are shown with a uniform intensity of 100; real data has noise, but the alignment-dependent behavior shown here does not depend on this simplification. Gray bars: the real microtubule structure. Squares below: the pixel grid our imaging samples, with the intensity value for each pixel. Target pixels (the ones being tested for gap) are highlighted — red when they cross the threshold (detected), amber when they don't (missed). Calculations below each scenario show local mean, threshold (local mean × 0.8), and whether the target pixel's intensity is below threshold.

### **Supplementary Tables**

#### **Supplementary Table 1. List of proteases tested with post-expansion staining in samples fixed with 3% w/v PFA and 0.1% w/v glutaraldehyde.**

HeLa cells were cultured on 12 mm glass coverslips in 24-well plates. Cells were fixed with freshly prepared 3% w/v PFA, 0.1% w/v GA in PBS for 10 min at room temperature (RT), followed by reduction with 0.1% w/v sodium borohydride in 1× PBS for 7 min at RT and then quenching with 100 mM glycine in 1× PBS for 10 min at RT. After fixation, cells were washed three times with 1× PBS for 5 min each at RT. To equip proteins with polymerizable handles, cells were then incubated with 0.1 mg/mL Acryloyl-X, SE (AcX) in PBS (diluted from a 10 mg/mL stock in anhydrous DMSO) for 2h at RT without shaking.

All image scale bars in the table are 50 µm (in physical units).

Protocol references are A: https://www.neb.com/en-us/protocols/2016/09/14/in-gel-digestion-protocol-for-endoproteinase-lysc-p8109 ; B: https://www.promega.com/resources/protocols/product-information-sheets/n/gluc-sequencing-grade-protocol/ ; C: https://www.promega.com/products/mass-spectrometry/peptidases-and-surfactants/lys-c-mass-spec-grade/ ; D: https://www.promega.com/products/mass-spectrometry/peptidases-and-surfactants/arg_c-sequencing-grade/ ; E: https://www.promega.com/products/mass-spectrometry/trypsin/trypsin-lys-c-mix-mass-spec-grade?catNum=V5071#protocols ; F: https://www.promega.com/products/mass-spectrometry/peptidases-and-surfactants/chymotrypsin-sequencing-grade/ .

| Enzyme(s) | Vendor | Cat# | Condition | Results |
| --- | --- | --- | --- | --- |
| Trypsin | New England Biolabs | P8101S | 50 ug/mL<br>50 mM Tris, pH 8<br>5 mM CaCl<sub>2</sub><br>37°C, 16h<br>Modified from ref. <sup>2</sup> | Microtubules appeared discontinuous<br>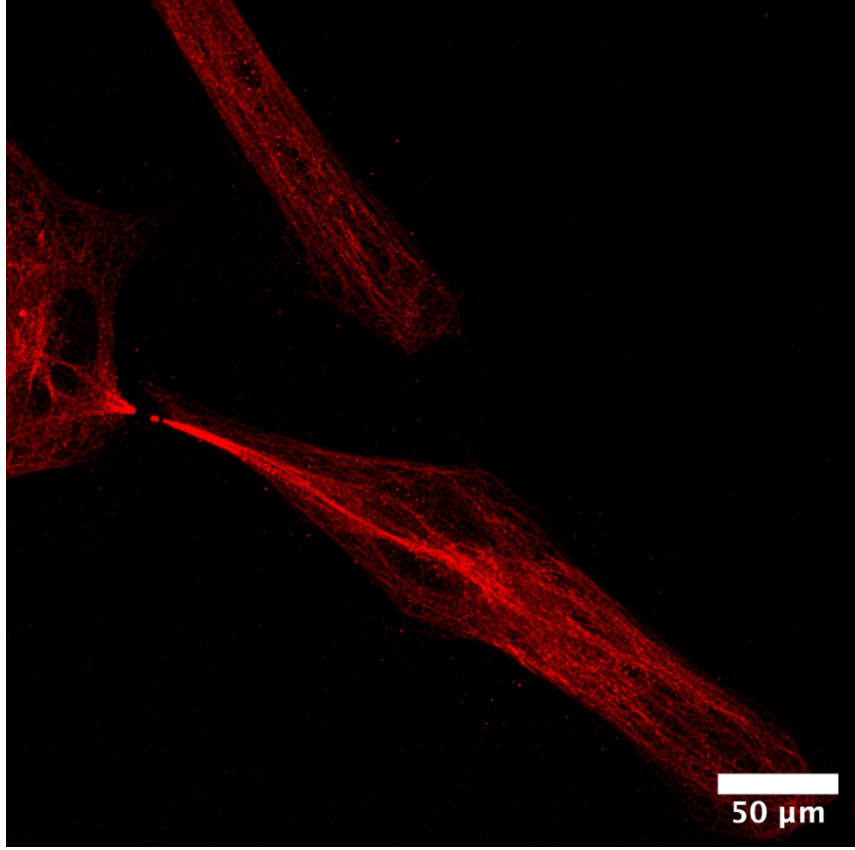<br>Brightness and contrast settings: first set by the ImageJ auto-scaling function and then manually adjusted to improve contrast for the stained structures of interest (done throughout the images in Supp. Table 1) |
| LysC | New England Biolabs | P8109S | 20 ug/mL<br>50 mM Tris, pH 8.5<br>1 mM EDTA<br>37°C, 16h<br>Modified from protocol reference A. | Microtubules appeared discontinuous<br>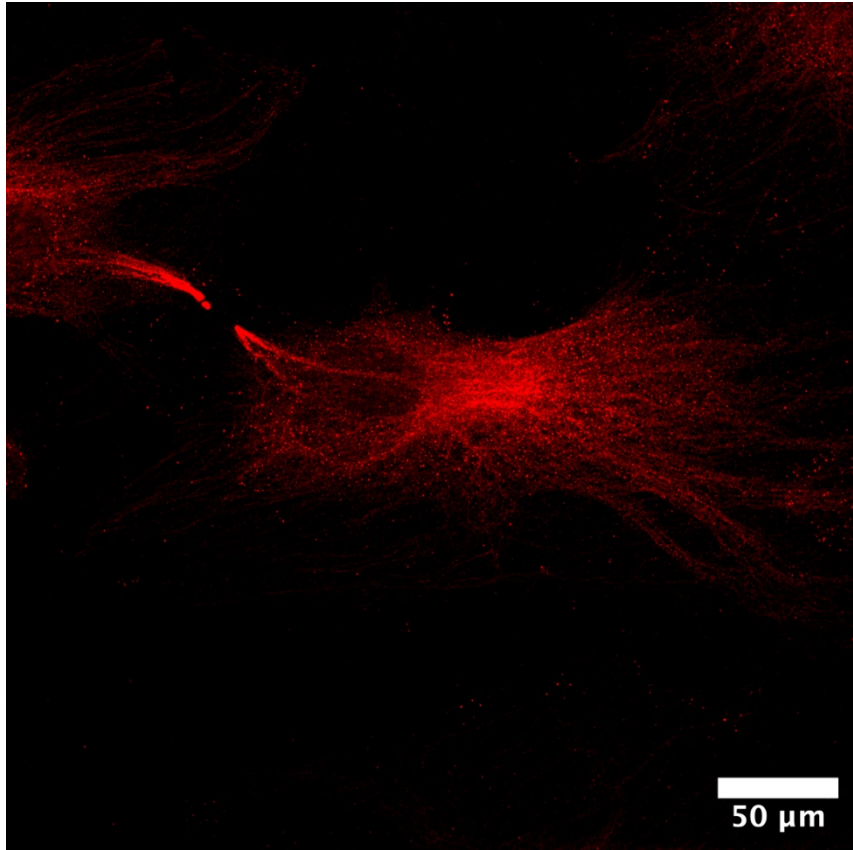 |
| GluC | New England Biolabs | P8100S | 50 ug/mL<br>100 mM ammonium bicarbonate buffer, pH 8<br>0.5 mM Glu-Glu<br>25°C, 16h | Cracks were observed within the gel & microtubules appeared discontinuous |
|  |  |  | Modified from protocol reference B. | 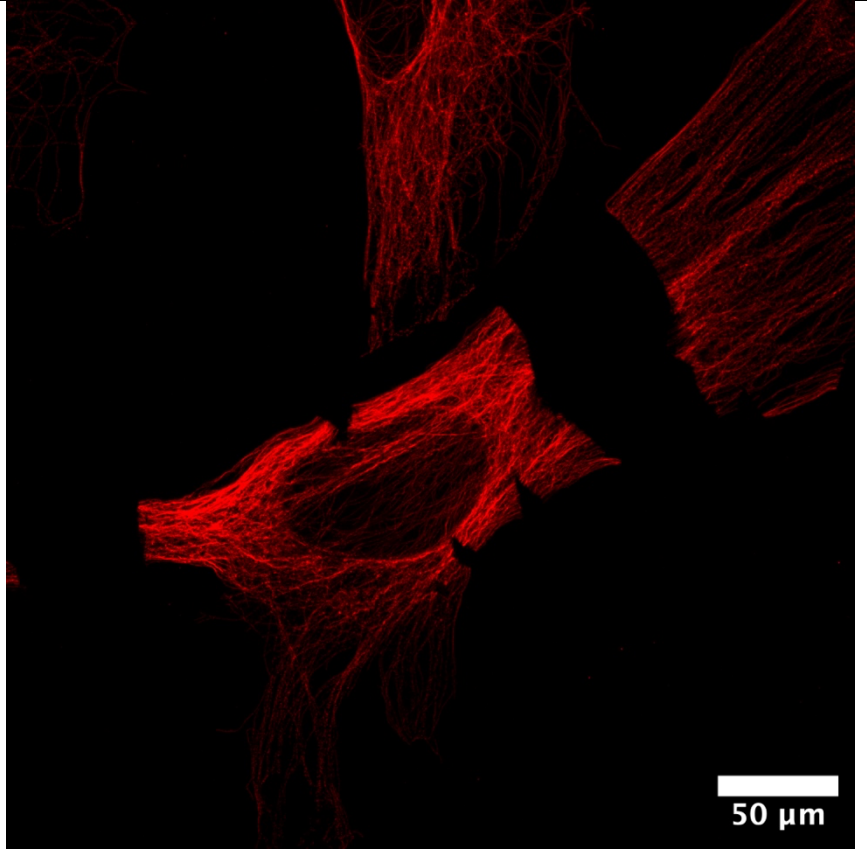 |
| AspN | New England Biolabs | P8104S | 50 ug/mL<br>50 mM ammonium bicarbonate buffer, pH 8<br>2.5mM ZnSO<sub>4</sub><br>37°C, 16h<br>Modified from protocol reference B. | Microtubules appeared discontinuous |
|  |  |  |  | 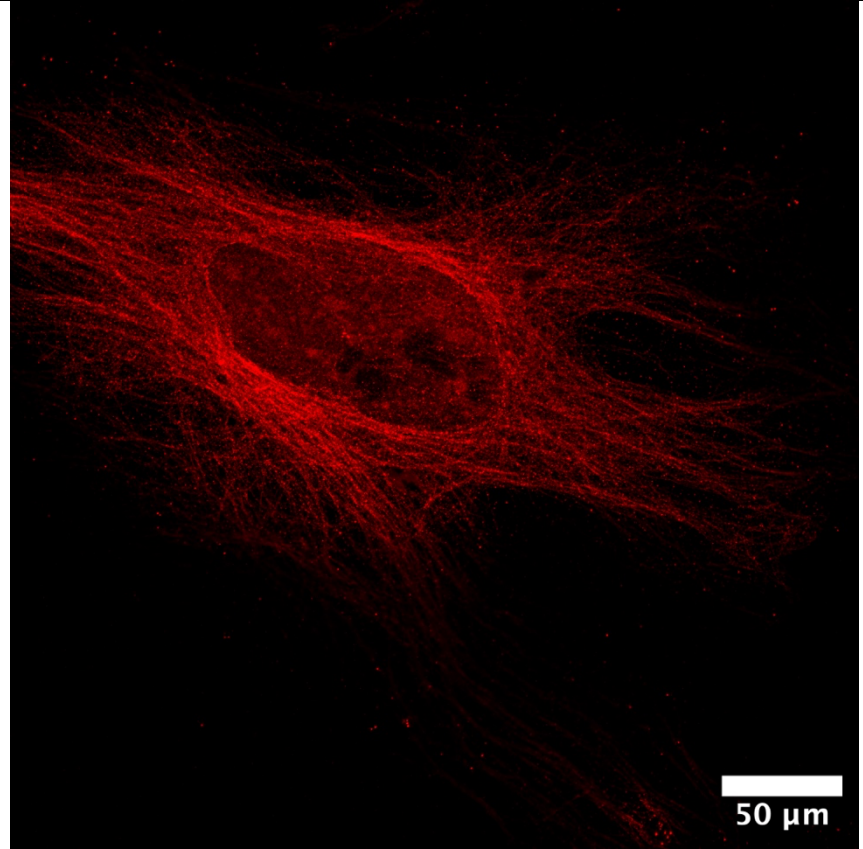 |
| rLys-C | Promega | V1671 | 15 ug/mL<br>50 mM Tris, pH 8.5<br>1 mM EDTA<br>37°C, 16h<br>Modified from protocol<br>reference C. | Microtubules appeared discontinuous |
|  |  |  |  | 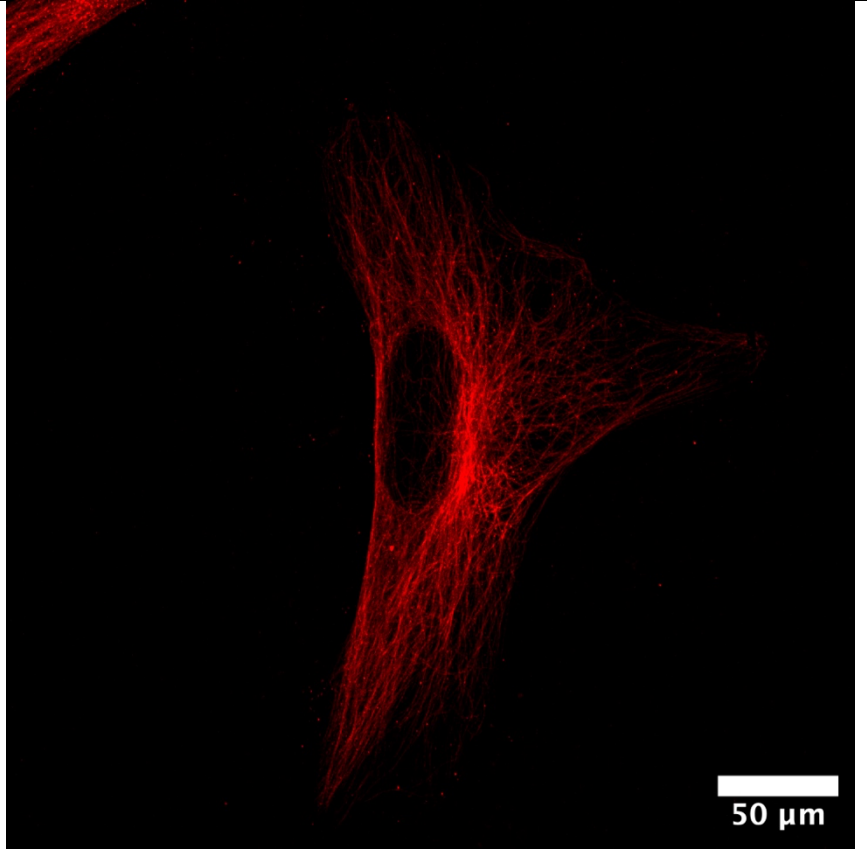 |
| Arg-C | Promega | V1881 | 10 ug/mL<br>50 mM Tris, pH 7.8<br>2 mM EDTA, 10 mM DTT<br>37°C, 16h<br>Modified from protocol<br>reference D. | Microtubules appeared discontinuous |
|  |  |  |  | 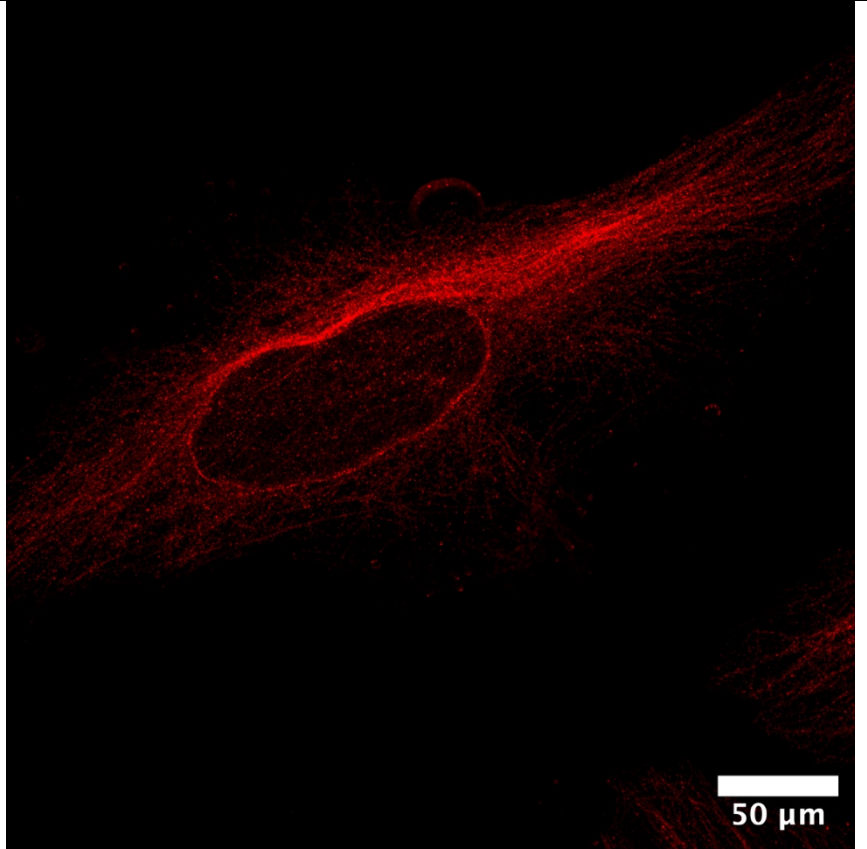 |
| LysN | Promega | VA1180 | 20 ug/mL<br>50 mM Tris, pH 8<br>2 mM ZnSO<sub>4</sub><br>37°C, 16h | Cracks were observed within the gel & microtubules appeared discontinuous |
|  |  |  |  | 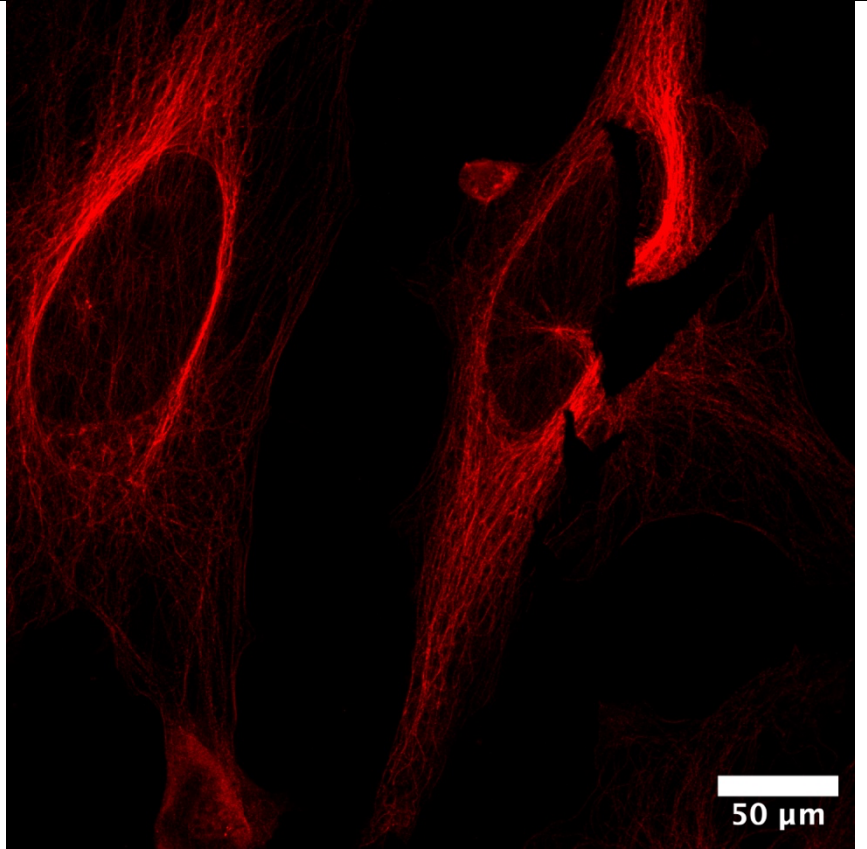 |
| LysC/Trypsin | Promega | V5071 | 20 ug/mL<br>50 mM Tris, pH 8<br>37°C, 2h<br>Modified from protocol<br>reference E. | Microtubules appeared discontinuous |
|  |  |  |  | 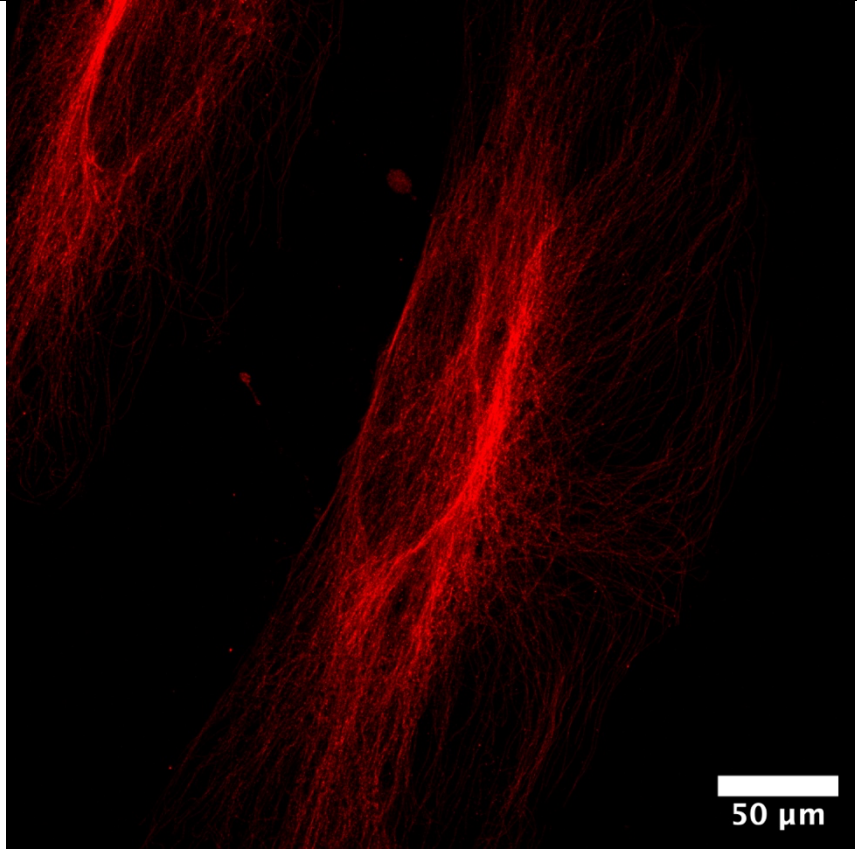 |
| Chymotrypsin | Promega | V1061 | 25 ug/mL<br>100 mM Tris, pH 8<br>25°C, 16h<br>Modified from protocol<br>reference F. | Cracks were observed within the gel & microtubules appeared discontinuous |

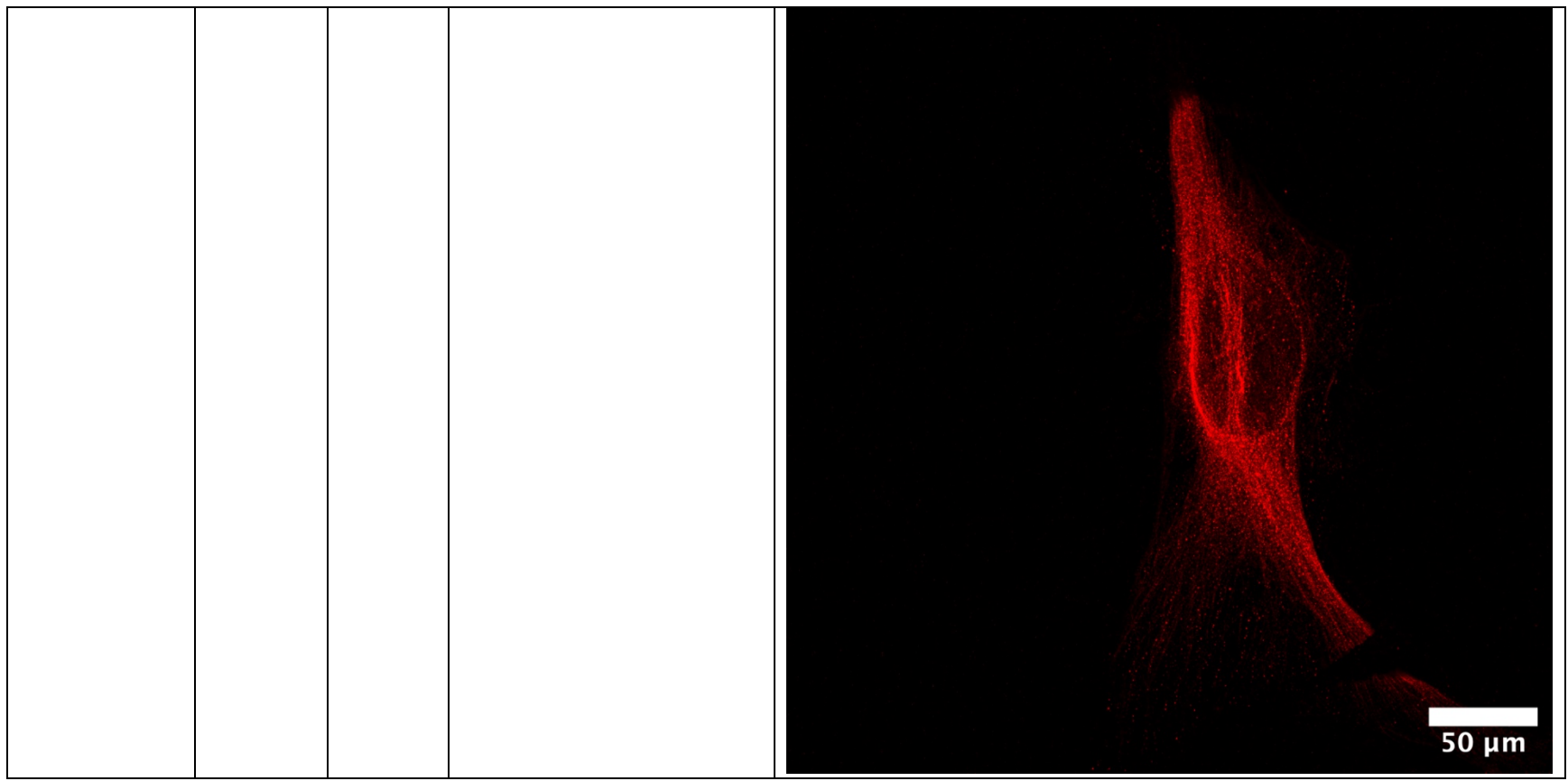

### Supplementary Table 2. Cleavable fixatives tested, and their outcomes.

**Design Principles.** We tried both custom-designed and commercially available cleavable crosslinking fixatives, guided by the following principles: first, the fixative molecule should be small enough to rapidly diffuse throughout the sample; second, the fixative should have multiple reactive groups that target functional groups on proteins (to achieve crosslinking); third, the fixative should be fast-acting to quickly lock biomolecules in place; and fourth, the crosslink should be cleavable. For the fixatives tested, this cleavability is afforded in one of three ways: (1) the cleavable bond is within the crosslinker itself and can later be broken by a cleavage reaction; (2) the cleavable bond is formed post-crosslinking and can later be broken by a cleavage reaction; or (3) the cleavable bond is formed at the conjugation between the crosslinker and the protein rather than in the crosslinker backbone itself.

| Vendor | Cat.# | Name | Chemical Structure | Cleavage | Results | Fixation results only, no ExM processing<br>(The lower panel is a magnified crop from the corresponding upper image) |
| --- | --- | --- | --- | --- | --- | --- |
| n.a.<br>(custom designed) | n.a.<br>(custom designed) | A1<br>(Sulfonyl-bis-acetaldehyde) | 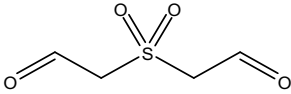 | Bears a sulfone bond which can be cleaved by sodium hydroxide | Crosslinking degree: 86%<br>(When 1.1% w/v A1 reacted with 100 mg/mL purified α-Lactalbumin, the proteins became a white precipitate, therefore unable to run SDS-PAGE gel to assess the crosslinking degree. Therefore, we used 10 mg/mL protein for reaction.)<br>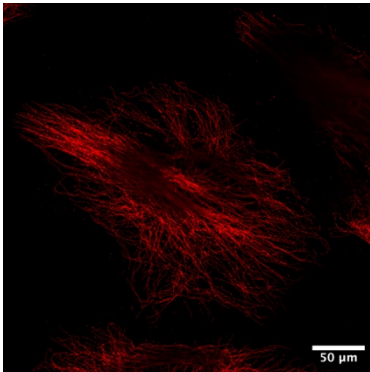 | 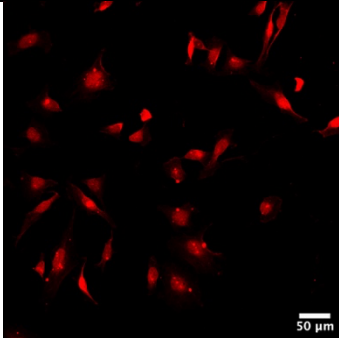<br>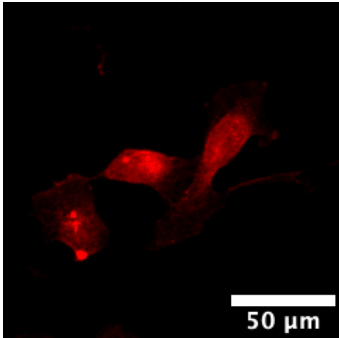<br>HeLa cells fixed with 4% v/v A1 in PBS, incubated for 6h at RT (fixation only, no ExM), and stained against beta-tubulin. The |
|  |  |  |  |  | 4% v/v A1 supplemented with 1% w/v PFA in PBS, 8h at RT, then post-expansion stained against beta-tubulin. The confocal image was acquired using a 40× water-immersion objective. Brightness and contrast settings: first set by the ImageJ auto-scaling function and then manually adjusted to improve contrast for the stained structures of interest (same for all the other images in Supp Table 2, except images without raw data)<br>Microtubules appear discontinuous after expansion | confocal image was acquired using a 10× air objective.<br>The cytoskeletal structure was almost completely gone. |
| n.a.<br>(custom designed) | n.a.<br>(custom designed) | A2<br>(2,2'-disulfanediyldiacetaldehyde) | O=CCSSCC=O | Bears a disulfide bond that can be cleaved by reduction (e.g. dithiothreitol) | Crosslinking degree: 7%<br>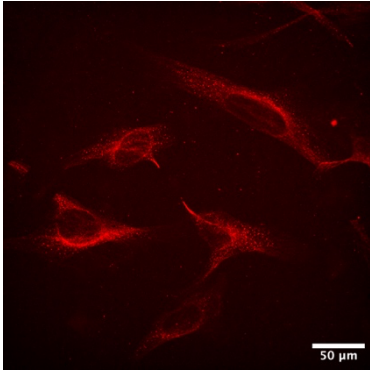<br>4% v/v A2 supplemented with 1% w/v PFA in PBS, 8h at RT, then stained against beta-tubulin. The confocal image was acquired using a 40× water-immersion objective.<br>Microtubules appear discontinuous | 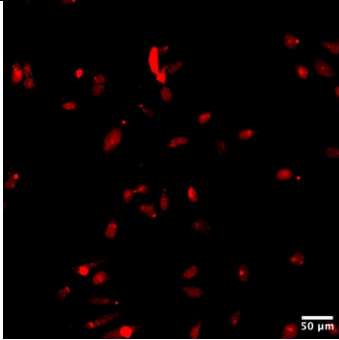 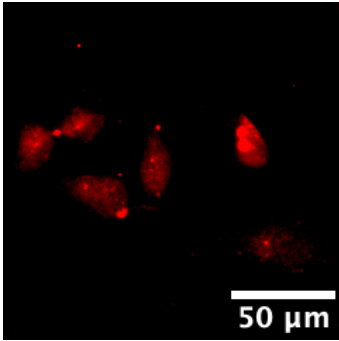<br>HeLa cells fixed with 4% v/v A2 in PBS, 6h at RT (fixation only, no ExM), and stained against beta-tubulin. The confocal image was acquired using a 10× air objective.<br>The cytoskeletal structure was almost completely gone. |
| n.a.<br>(custom<br>designed) | n.a.<br>(custom<br>designed) | CL1<br>(tartaric<br>aldehyde) | O=C[C@@H](O)[C@H](O)C=O | Bears a diol<br>bond that can<br>be cleaved<br>by oxidation<br>(e.g. sodium<br>periodate) | No crosslinking (PAGE gel<br>result) | 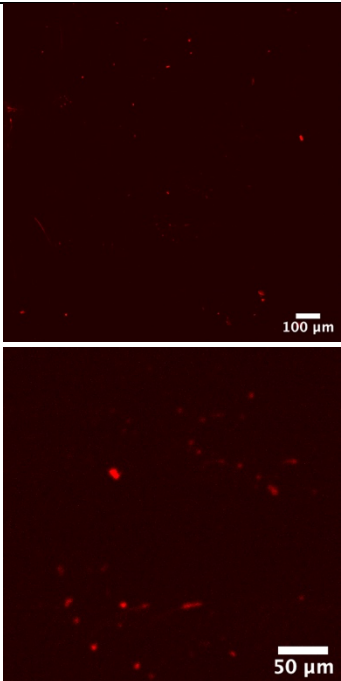<br>HeLa cells fixed with 4%<br>v/v CL1 in PBS, 6h at RT<br>(fixation only, no ExM),<br>and stained against beta-tubulin. The confocal<br>image was acquired<br>using a 10× air objective.<br>No tubulin signal detected |
| n.a.<br>(custom designed) | n.a.<br>(custom designed) | CL2<br>(bis(chlorocarbonyl)(carbamoyl)disulfane) | 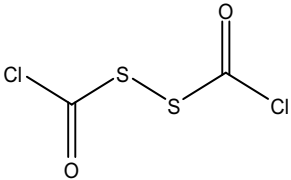 ClC(=O)SSC(=O)Cl | Bears a disulfide bond that can be cleaved by reduction | No crosslinking<br>(PAGE gel result) | 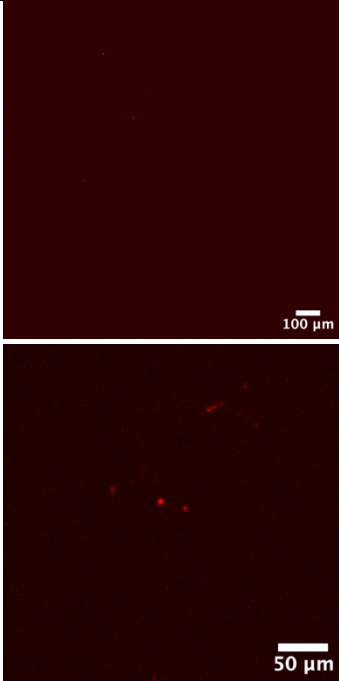<br>HeLa cells fixed with 4% v/v CL2 in PBS, 6h at RT (fixation only, no ExM), and stained against beta-tubulin. The confocal image was acquired using a 10× air objective.<br>No tubulin signal detected |
| n.a.<br>(custom designed) | n.a.<br>(custom designed) | B1<br>(2,2'-<br>(sulfon<br>ylbis(m<br>ethylen<br>e))bis(<br>oxiran<br>e)) | 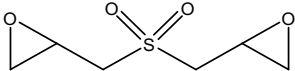 | Bears a sulfone bond which can be cleaved by sodium hydroxide | Crosslinking degree 4%<br>Efficient epoxy-based crosslinking generally requires alkaline conditions, as epoxide groups react efficiently with protein sulfhydryl, amine, and hydroxyl nucleophiles only at elevated pH<sup>10</sup>. At near-neutral pH, their reactivity toward protein nucleophiles is low and intermolecular reactions with proteins proceed very slowly<sup>11,12</sup>, so only long incubations substantially increase crosslinking in vitro, which limits their straightforward use in live-cell applications. | 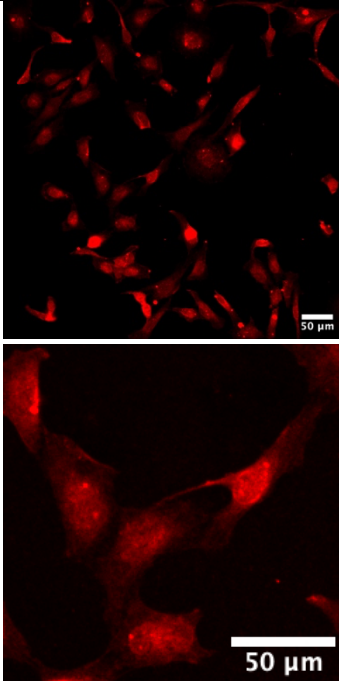<br>HeLa cells fixed with 4% v/v B1 in PBS, 6h at RT (fixation only, no ExM), and stained against beta-tubulin. The confocal image was acquired using a 10× air objective.<br>Tubulins were dispersed. |
| n.a.<br>(custom designed) | n.a.<br>(custom designed) | B2<br>(1,2-bis(oxiran-2-ylmethyl)disulfane) | 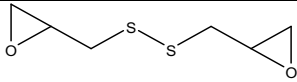 | Bears a disulfide bond that can be cleaved by reduction | Crosslinking degree = 11%<br>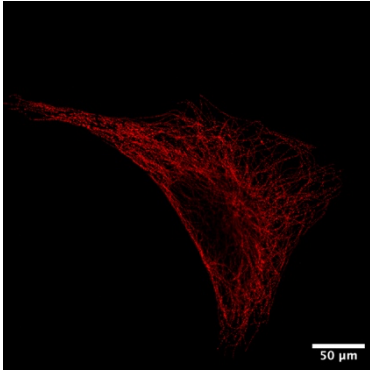<br>4% v/v B2 supplemented with 3% PFA in PBS, 8h at RT, then stained against beta-tubulin. The confocal image was acquired using a 40× water-immersion objective.<br>Microtubules appear discontinuous<br>Note: We aimed to minimize crosslinking between proteins; therefore, we initially tested 1% PFA as a starting point (the choice of 1% was arbitrary). However, preservation was | 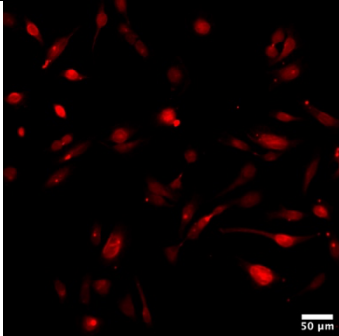 <br>HeLa cells fixed with 4% v/v B2 in PBS, 6h at RT (fixation only, no ExM), and stained against beta-tubulin. The confocal image was acquired using a 10× air objective.<br>The cytoskeletal structure was almost completely gone. |
|  |  |  |  |  | very poor (similar to the condition without PFA), so we increased the PFA concentration to 3%. The same applies to all subsequent conditions where 3% PFA was used instead of 1% PFA. |  |
| n.a.<br>(custom designed) | n.a.<br>(custom designed) | B3<br>(1,2-di(oxiran-2-yl)ethane-1,2-diol) |  | Bears a diol bond that can be cleaved by oxidation | Crosslinking degree: 4%<br>Note: For all crosslinkers with diol bonds (regardless of whether they are within the crosslinker or formed upon crosslinking), we qualitatively observed that after cleavage, the tubulin signal appears aggregated into clusters. This is likely because after cleavage, aldehyde groups are generated, which subsequently acts as new crosslinkers. This interpretation is consistent with the known periodate-mediated oxidation/cleavage of vicinal diols to aldehydes, which |  <br>HeLa cells fixed with 4% v/v B3 in PBS, 6h at RT (fixation only, no ExM), |
|  |  |  |  |  | can subsequently form additional covalent bonds with nucleophiles such as amines and are exploited in periodate–lysine–paraformaldehyde (PLP) fixation protocols <sup>13</sup>. | and stained against beta-tubulin. The confocal image was acquired using a 10× air objective.<br>Tubulins were dispersed. |
| n.a.<br>(custom designed) | n.a.<br>(custom designed) | Tracel ess crossli nker Bi-APB |  | The reaction between amines and boronic acids is reversible <sup>3</sup> | No crosslinking (PAGE gel result) | Not tried on cells |
| Sigma Aldrich | 202533 | DEB (1,3-Butadi ene diepoxi de) |  | Bears no cleavable bond but a cleavable diol bond will form upon crosslinking | Crosslinking degree: 6%<br><br>4% v/v DEB supplemented with 1% w/v PFA in PBS, 8h at RT, then stained against beta-tubulin post-expansion staining. The confocal |  |
|  |  |  |  |  | image was acquired using a 40× water-immersion objective.<br>Microtubules appear discontinuous after expansion, and tubulin seems aggregated in some areas, possibly because after diol cleavage, new aldehyde groups are generated, leading to protein crosslinking and aggregation. | <br>HeLa cells fixed with 4% v/v DEB in PBS, 6h at RT (fixation only, no ExM), and stained against beta-tubulin<br>Tubulins were dispersed. |
| ThermoFisher | 22585 | DSP (dithio bis(succinimidyl propionate)) |  | Bears a disulfide bond that can be cleaved by reduction | Crosslinking degree: 62 %<br><br>12.5 mM DSP supplemented with 1% w/v |  |
|  |  |  |  |  | PFA in PBS, 8h at RT, then stained against beta-tubulin. The confocal image was acquired using a 40× water-immersion objective.<br>Microtubules appear discontinuous and have cracks within the structure. | <br>HeLa cells fixed with 20 mM DSP in phosphate buffer (pH 8), 6 h at RT (fixation only, no ExM), and stained against beta-tubulin.<br>Tubulins were dispersed. |
| ThermoFisher | 20589 | DST<br>(disuccinimidyl tartrate) |  | Bears a diol bond that can be cleaved by oxidation | Crosslinking degree: 50% |  |
|  |  |  |  |  |  | <br>HeLa cells were fixed with 20 mM DST in PBS for 6 hours at RT, then stained against beta-tubulin. The confocal image was acquired using a 10× air objective.<br>Tubulins were dispersed.<br>Raw confocal files were unavailable; therefore, scale bars are not shown. |
| ThermoFisher | 21600 | BSOC OES (bis[2-(succinimidyl)oxy]carbonylox) |  | Bears a sulfone bond which can be cleaved by sodium hydroxide | Crosslinking degree: 75 % |  |
|  |  | y)ethyl]<br>sulfone<br>) |  |  | <br>12.5 mM of BSOCOES supplemented with 1% w/v PFA in PBS, 8h at RT, then post expansion stained against beta-tubulin. The confocal image was acquired using a 40× water-immersion objective.<br>Microtubules appear discontinuous |  |
| ThermoFisher | 21565 | EGS<br>(ethylene glycol bis(succinimidyl |  | Bears two ester sites can be cleaved with hydroxylamine | Crosslinking degree was not recorded<br> |  |
|  |  | succinate)) |  |  |  | <br>HeLa cells were fixed with 6.25 mM EGS + 1% PFA, in PBS, incubated for 6h at RT (fixation only, no ExM), and stained against beta tubulin. The confocal image was acquired using a 40× water-immersion objective.<br>Tubulins were dispersed. |

|  |  |  |  |  |  |  |
| --- | --- | --- | --- | --- | --- | --- |
|  |  |  |  |  |  | Raw confocal files were unavailable; therefore, scale bars are not shown. |
| ThermoFisher | 21578 | Sulfo-DSP (3,3'-dithiobis(sulfo succinimidyl propionate)) |  | Bears a disulfide bond that can be cleaved by reduction | Crosslinking degree: 60% | We did not test this in cells because sulfo-DSP also bears a disulfide bond, like DSP, and we found that the cleavage efficiency of DSP was not high enough to give us a crack-free gel. |
| ThermoFisher | 26169 | SDAD (NHS-SS-Diazirine) |  | Bears a disulfide bond that can be cleaved by reduction | Crosslinking degree: 73% | Not evaluated in cells. At the time of the initial screen, we had not established whether the photoactivation required for SDAD-based fixation would preserve cellular biomolecules, including nucleic acids. We therefore did not advance these candidates to cellular testing. During subsequent development of nfExM, we adopted a brief, UV exposure for the separate purpose of initiating rapid hydrogel |
| | | | | | | polymerization at $-20^{\circ}\text{C}$, where APS/TEMED was ineffective. Photoinitiated gelation has previously been demonstrated in PhotoExM<sup>14</sup>. The effects of the nfExM irradiation condition on DNA and RNA integrity were not assessed. |
| ThermoFisher | 26175 | Sulfo-SDAD (Sulfo-NHS-SS-Diazirine) |  | Bears a disulfide bond that can be cleaved by reduction | Crosslinking degree: 66% | Not evaluated in cells. See previous row for reasoning. |
| ProteoChem | c1116 | SPDP (succinimidyl 3-(2-pyridyl dithio) propionate) |  | Bears a disulfide bond that can be cleaved by reduction | Crosslinking degree: 44% | We did not test this in cells because SPDP also bears a disulfide bond, like DSP, and we found that the cleavage efficiency of DSP was not high enough to give us a crack-free gel. |
| ThermoFisher | 20665 | DTBP<br>(Dimethyl 3,3'-dithiobispropionimidate•2HCl) | COC(=O)CC[NH2+][Cl-]SSCC(=O)N[Cl-] | Bears a disulfide bond that can be cleaved by reduction | Crosslinking degree: 73%<br><br>1.25 mM DTBP + 3% PFA, in PBS for 8h at RT, post-expansion stained against beta tubulin. The confocal image was acquired using a 40× water-immersion objective. | <br>HeLa cells were fixed with 20mM DTBP in phosphate buffer (pH 8) incubated for 1h (without PFA, without ExM). The confocal image was acquired using a 40× water-immersion objective.<br>Tubulins were dispersed |
| ThermoFisher | 21666 | DMP<br>(Dimethyl pimelidate •2 HCl) |  | Bis(imidoester) crosslinkers form amidine bonds with primary amines, which have been reported to undergo hydrolysis at high pH, rendering the crosslinks reversible at high pH. In this study, we used DMA, DMP and DMS for in vitro crosslinking and did not test any high-pH cleavage protocol. | Crosslinking degree: 39% | <br>HeLa cells were fixed with 6.25 mM DMP + 1% PFA, in PBS, incubated for 6h at RT (fixation only, no ExM), and stained against beta tubulin. The confocal image was acquired using a 10× air objective.<br>Tubulins were dispersed. |
|  |  |  |  |  |  | Raw confocal files were unavailable; therefore, scale bars are not shown. |
| ThermoFisher | 20660 | DMA (Dimethyl adipimide•2 HCl) | COC(=O)C(CCCCNC(=O)OC)C(=O)OC |  | Crosslinking degree: 26% | <br>HeLa cells were fixed with 6.25 mM DMA + 1% PFA, in PBS, incubated for 6h at RT (fixation only, no ExM), and stained against beta tubulin. The confocal image was |

|  |  |  |  |  |  |
| --- | --- | --- | --- | --- | --- |
|  |  |  |  |  | acquired using a 10× air objective.<br>Tubulins were dispersed.<br>Raw confocal files were unavailable; therefore, scale bars are not shown. |
| ThermoFisher | 20700 | DMS (Dimethylsuberimide •2 HCl) | COC(=O)CCCCCCCCC(=O)N[CH2+][Cl-] | Crosslinking degree: 37% | <br>HeLa cells were fixed with 6.25 mM DMS + 1% PFA, in PBS, incubated |

|  |  |  |  |  |  |  |
| --- | --- | --- | --- | --- | --- | --- |
|  |  |  |  |  |  | for 6h at RT (fixation only, no ExM), and stained against beta tubulin. The confocal image was acquired using a 10× air objective.<br>Tubulins were dispersed<br>Raw confocal files were unavailable; therefore, scale bars are not shown. |
| ThermoFisher | 21525 | DFDN B (1,5-difluoro-2,4-dinitrobenzene) |  | DFDNB contains aryl halide groups that can form cross-links with protein sulfhydryl groups and have been reported to be reversible upon treatment with excess thiol (e.g., DTT) (manufacture | No crosslinking (PAGE gel result) |  |
|  |  |  |  | r's instructions and ref. <sup>15</sup>). In this study, we only used DFDNB for crosslinking and did not test the cleavage step. |  | <br>HeLa cells were fixed with 6.25 mM DFDNB + 1% PFA in PBS, incubated for 6h at RT (fixation only, no ExM), and stained against beta tubulin. The confocal image was acquired using a 10× air objective.<br>The cytoskeletal structure was almost completely gone. |

**Supplementary Table 3: Detailed Statistics for Figure 2**

| Metric | Method | n | Mean ± SD | Median (IQR) | 95% CI |
| --- | --- | --- | --- | --- | --- |
| Mean Intensity (a.u.) | proExM(PFA) | 75 | 42.74 ± 11.85 | 43.71 (33.12–50.59) | (40.01, 45.46) |
| Mean Intensity (a.u.) | CryoExM | 124 | 76.24 ± 41.67 | 68.42 (49.71–98.81) | (68.83, 83.64) |
| Mean Intensity (a.u.) | nfExM | 155 | 291.48 ± 88.03 | 284.60 (228.75–338.83) | (277.51, 305.45) |
| Gap Ratio (%) | proExM(PFA) | 75 | 28.27 ± 5.20 | 28.57 (24.64–31.93) | (27.07, 29.46) |
| Gap Ratio (%) | CryoExM | 124 | 24.17 ± 10.06 | 24.29 (16.67–30.16) | (22.38, 25.96) |
| Gap Ratio (%) | nfExM | 155 | 1.80 ± 2.07 | 1.30 (0.00–2.93) | (1.48, 2.13) |
| Gap Size (μm) | proExM(PFA) | 75 | 0.09 ± 0.02 | 0.09 (0.08–0.10) | (0.09, 0.10) |
| Gap Size (μm) | CryoExM | 124 | 0.09 ± 0.02 | 0.09 (0.07–0.10) | (0.09, 0.09) |
| Gap Size (μm) | nfExM | 155 | 0.03 ± 0.03 | 0.04 (0.00–0.04) | (0.03, 0.04) |

**Statistical comparisons (Mann–Whitney U test):**

| Metric | Comparison | P value | Significance |
| --- | --- | --- | --- |
| Mean Intensity (a.u.) | proExM(PFA) vs CryoExM | 5.84e-12 | **** |
| Mean Intensity (a.u.) | proExM(PFA) vs nfExM | 1.08e-34 | **** |
| Mean Intensity (a.u.) | CryoExM vs nfExM | 8.76e-46 | **** |
| Gap Ratio (%) | proExM(PFA) vs CryoExM | 3.35e-04 | **** |
| Gap Ratio (%) | proExM(PFA) vs nfExM | 2.92e-35 | **** |
| Gap Ratio (%) | CryoExM vs nfExM | 1.74e-46 | **** |
| Gap Size (μm) | proExM(PFA) vs CryoExM | 0.050 | ns |
| Gap Size (μm) | proExM(PFA) vs nfExM | 1.80e-31 | **** |
| Gap Size (μm) | CryoExM vs nfExM | 1.02e-38 | **** |

Significance levels: n.s. not significant, \* $p < 0.05$, \*\* $p < 0.01$, \*\*\* $p < 0.001$, \*\*\*\* $p < 0.0001$

**Supplementary Table 4: Detailed Statistics for Figure 3**

| Method | Intensity (a.u., Mean ± SD) | Gap Ratio (% , Mean ± SD) | Gap Size (μm, Mean ± SD) | N segments |
| --- | --- | --- | --- | --- |
| high T% ProExM | 378.99 ± 95.62 | 11.58 ± 5.85 | 0.074 ± 0.029 | 59 |
| nfExM | 912.38 ± 144.47 | 1.65 ± 2.28 | 0.029 ± 0.032 | 61 |

**Statistical comparisons (Mann-Whitney U-test):**

| Comparison | Parameter | p-value | Significance |
| --- | --- | --- | --- |
| nfExM vs. high T% ProExM | Intensity | $3.60 \times 10^{-21}$ | **** |
| high T% ProExM vs. nfExM | Gap Ratio | $1.52 \times 10^{-16}$ | **** |
| high T% ProExM vs. nfExM | Gap Size | $2.49 \times 10^{-11}$ | **** |

Significance levels: \*\*\*\* $p < 0.0001$

**Supplementary Table 5: Detailed Statistics for Figure 4**

| Method | Intensity (a.u., Mean ± SD) | Gap Ratio (% Mean ± SD) | Gap Size (μm, Mean ± SD) | N segments |
| --- | --- | --- | --- | --- |
| ProExM | 503.64 ± 153.84 | 3.07 ± 3.55 | 0.038 ± 0.028 | 174 |
| ExPath | 502.38 ± 128.45 | 2.45 ± 3.01 | 0.037 ± 0.026 | 193 |
| MAP | 623.35 ± 201.87 | 2.75 ± 2.95 | 0.038 ± 0.032 | 145 |
| nfExM | 1029.47 ± 200.56 | 1.26 ± 1.88 | 0.029 ± 0.032 | 190 |

**Statistical comparisons (Mann-Whitney U-test):**

| Comparison | Parameter | p-value | Significance |
| --- | --- | --- | --- |
| nfExM vs. ProExM | Intensity | $5.68 \times 10^{-55}$ | **** |
| nfExM vs. ExPath | Intensity | $2.08 \times 10^{-60}$ | **** |
| nfExM vs. MAP | Intensity | $1.68 \times 10^{-38}$ | **** |
| ProExM vs. ExPath | Intensity | 1.000 | n.s. |
| ProExM vs. MAP | Intensity | $3.04 \times 10^{-7}$ | **** |
| ExPath vs. MAP | Intensity | $1.88 \times 10^{-7}$ | **** |
| ProExM vs. nfExM | Gap Ratio | $1.57 \times 10^{-7}$ | **** |
| ExPath vs. nfExM | Gap Ratio | $1.18 \times 10^{-5}$ | **** |
| MAP vs. nfExM | Gap Ratio | $9.15 \times 10^{-7}$ | **** |
| ProExM vs. ExPath | Gap Ratio | 1.000 | n.s. |
| ProExM vs. MAP | Gap Ratio | 1.000 | n.s. |
| ExPath vs. MAP | Gap Ratio | 1.000 | n.s. |
| ProExM vs. nfExM | Gap Size | $9.80 \times 10^{-4}$ | *** |
| ExPath vs. nfExM | Gap Size | 0.004 | ** |
| MAP vs. nfExM | Gap Size | 0.017 | * |
| ProExM vs. ExPath | Gap Size | 1.000 | n.s. |
| ProExM vs. MAP | Gap Size | 1.000 | n.s. |
| ExPath vs. MAP | Gap Size | 1.000 | n.s. |

P-values are Bonferroni-corrected for 6 pairwise comparisons (4 methods)

Significance levels: n.s. not significant, \* $p < 0.05$, \*\* $p < 0.01$, \*\*\* $p < 0.001$, \*\*\*\* $p < 0.0001$.

**Supplementary Table 6: Detailed Statistics for Figure 5**

| Metric | Method | n | Mean ± SD | Median (IQR) | 95% CI |
| --- | --- | --- | --- | --- | --- |
| GM130<br>Signal Pixel<br>Counts | CryoExM | 16 | 42401.3 ±<br>29266.6 | 31563.5<br>(25747.8–<br>52116.0) | (26806.3,<br>57996.4) |
| GM130 Mean<br>Intensity<br>(a.u.) | CryoExM | 16 | 135.7 ± 19.4 | 137.9 (123.3–<br>146.3) | (125.4, 146.0) |
| GM130<br>Signal Pixel<br>Counts | nfExM | 16 | 1355002.3 ±<br>457558.6 | 1322288.0<br>(1182253.5–<br>1500040.0) | (1111186.5,<br>1598818.1) |
| GM130 Mean<br>Intensity<br>(a.u.) | nfExM | 16 | 256.7 ± 32.1 | 255.9 (232.7–<br>272.9) | (239.6, 273.8) |
| TOM20<br>Signal Pixel<br>Counts | CryoExM | 8 | 652449.8 ±<br>144015.6 | 640159.5<br>(559697.5–<br>770053.0) | (532049.7,<br>772849.8) |
| TOM20<br>Mean<br>Intensity<br>(a.u.) | CryoExM | 8 | 269.7 ± 47.3 | 258.1 (236.0–<br>280.8) | (230.1, 309.3) |
| TOM20<br>Signal Pixel<br>Counts | nfExM | 8 | 1117785.2 ±<br>289200.7 | 1099425.0<br>(959146.8–<br>1208696.5) | (876007.4,<br>1359563.1) |
| TOM20<br>Mean<br>Intensity<br>(a.u.) | nfExM | 8 | 421.7 ± 48.9 | 414.9 (383.5–<br>437.0) | (380.8, 462.6) |

**Statistical comparisons (Unpaired t-test):**

| Metric | Comparison | P value | Significance |
| --- | --- | --- | --- |
| GM130 Signal Pixel<br>Counts | nfExM vs CryoExM | 1.54e-06 | **** |
| GM130 Mean Intensity<br>(a.u.) | nfExM vs CryoExM | 1.54e-06 | **** |
| TOM20 Signal Pixel<br>Counts | nfExM vs CryoExM | 0.0011 | *** |
| TOM20 Mean Intensity<br>(a.u.) | nfExM vs CryoExM | 1.55e-04 | **** |

Significance levels: n.s. not significant, \* $p < 0.05$, \*\* $p < 0.01$, \*\*\* $p < 0.001$, \*\*\*\* $p < 0.0001$.
